# Metabolic task analysis reveals distinct metabotypes in end-stage dilated cardiomyopathy

**DOI:** 10.1101/2025.03.28.645901

**Authors:** Bastien S.C. Nihant, Job A.J. Verdonschot, Sonia Bălan, Eva Thielecke, Joost J.F.P Luiken, Miranda Nabben, Stephane Heymans, Marian Breuer, Michiel E. Adriaens

## Abstract

Dilated cardiomyopathy (DCM) is associated with shifts in cardiac metabolism. Those shifts are inconsistent between patients, possibly due to heterogeneity in DCM etiologies. Identifying metabolic subtypes, or metabotypes, in DCM patients may open personalized treatment opportunities. Developing a methodology to identify metabotypes would be a boon in this regard. Here, we describe a metabotyping pipeline, integrating advanced metabolic modeling methods optimized for cardiac research, to uncover these subtypes using widely available transcriptomics data. We applied our method to publicly available cardiac data of end-stage DCM patients and non-failing controls, identifying two metabotypes in the DCM group. These metabotypes are characterized by unique metabolic alterations, notably in calcium handling, amino-acid oxidation, and the pentose phosphate pathway. Strikingly, one metabotype exhibited a greater deviation from healthy controls, suggesting a greater metabolic contribution to its underlying etiology. Further transcriptome-wide analysis revealed immune-related differences between metabotypes, suggesting an interplay between inflammation, immune response and metabolism in these DCM subtypes. Our study uncovers cardiometabolic heterogeneity in DCM and underscores the potential of transcriptome-derived metabotyping in cardiovascular research.

## Introduction

The heart is one of the most energy-demanding organs in the human body, estimated to consume 400 kcal per day, or 15-20% of total caloric usage (1). To sustain its constant activity the heart has evolved a metabolism capable of utilizing various nutrients, a concept known as metabolic flexibility. Fatty acids in particular represent about 60% of substrate usage in the healthy heart (2). Dysregulations of cardiac substrate usage have been observed in heart failure (HF). More specifically, cardiac metabolism may shift away from fatty acid oxidation and towards glycolysis (3). Such a shift has also been observed in dilated cardiomyopathy (DCM), a non-ischemic form of HF (4). These metabolic changes, however, have recently been under some scrutiny. Targeting metabolism as a treatment strategy has proven to be difficult (5). While, for example, sodium-glucose cotransporter-2 inhibitors (SGLT2i) have been shown to be beneficial in DCM treatment, their effect is much broader than glucose metabolism and it is still unclear through which mechanisms they benefit DCM patients (6). Furthermore, recent studies have challenged the extent of loss of metabolic flexibility in non-ischemic HF (7). This underlines the need for further research in this field to provide more definitive answers.

An important consideration is that DCM is largely thought of as a phenotype arising from various genetic and non-genetic disease triggers. Even prevalent DCM-related variants, such as truncating Titin variants, require environmental factors to trigger the phenotype and many cases are still idiopathic (8). For this reason, it is also accepted that the molecular profile of the DCM heart may vary between patients. Previous studies showed distinct transcriptomic and metabolic profiles correlating with truncated titin variants and with disease severity in non-genetic DCM patients (9,10). It is therefore reasonable to postulate that cardiac metabolism may be impacted differently in different subjects irrespective of severity, which may explain the lack of success of broadly metabolism-oriented treatments (5). Such differences between metabolic profiles of patients are often referred to as metabotypes. This concept is particularly useful in personalized nutrition and medicine when tailoring treatment and dietary recommendations to individuals, thus avoiding one-size-fits-all approaches (11). Different underlying metabotypes could in part explain the heterogeneity in results from studies of DCM cardiac metabolism.

Studying metabolic changes in human subjects is, however, a complicated and costly exercise (12). The gold standard method for quantifying metabolic changes is to measure metabolic fluxes, using isotopic tracers such as 2H and 13C followed by mass spectrometry. Such 2H/13C flux analysis reliably measures the metabolic conversions by following the fate of the ingested or injected isotopes (13). Recent advances have enabled the use of magnetic resonance spectroscopy (MRS) to infer metabolic rates, enabling non-invasive studies of specific metabolites in living patients (14).

These methods are invaluable to investigate existing hypotheses. However, in some cases, being able to rapidly and cheaply generate insights from widely available data would be of significant value. For this reason, systems biology approaches to generate and refine hypotheses that could be verified using gold-standard methods are invaluable. Genome-scale metabolic models (GEMs) are one such approach (15). GEMs consist of a network of reactions and metabolites, aiming at complete coverage of the metabolism of an organism with accurate stoichiometry. Reactions are modelled using steady-state fluxes representing the homeostatic reaction activity over time. Such steady-state flux-based modelling therefore simplifies the dynamics of metabolic responses over time in exchange for allowing to model the complete metabolism. Multiple types of methods exist to derive subject-specific insights from these generalized models.

In this paper we refined an approach to directly infer metabolic functionality by combining a comprehensive human GEM and subject level omics data (16). This approach is referred to as metabolic task analysis. Metabolic tasks describe net conversions of biological interest, for example the oxidation of glucose with oxygen to CO_2_ and water under production of ATP; metabolic tasks are defined in terms of their substrates’ and products’ stoichiometries. The metabolic task activity is then evaluated from the expression of the enzymes catalyzing the reaction pathways from substrates to products. In this way, metabolic tasks allow to query the transcriptomics data for specific metabolic functions (17). This is particularly valuable, as it unlocks new metabolic insights specifically from existing transcriptomics data, one of the most readily available and widely utilized data types in molecular cardiology.

The generated task activity scores represent a detailed metabolic fingerprint and can be used for group comparisons and, importantly, metabotyping. Metabolic task analysis offers important advantages over traditional methods such as pathway enrichment analysis. Pathway enrichment analysis only indicates whether a pathway contains more differentially expressed genes than expected by chance. How to subsequently interpret the activity of such a pathway is not straightforward. In contrast, metabolic task analysis identifies bottlenecks in such pathways and provides direct activity scores with a clear biological implication. Current implementations of metabolic task analysis, however, have limitations in their applicability to cardiac transcriptomic data. Namely, these tools are based on older GEMs and do not offer tasks related to myosin contraction or calcium handling in the sarcoplasmic reticulum (SR), hampering their application to the study of cardiac metabolism (16,18,19).

Here we present a metabotyping pipeline specific for the heart in health and disease. We use the most recent human GEM and a list of metabolic tasks, both extended to cover cardiac-specific functionalities (16,18,20). We then use our pipeline on publicly available cardiac transcriptomic data from DCM subjects and healthy controls, with the hypothesis that distinct metabotypes would be apparent in DCM subjects. We do not only uncover distinct metabotypes, but also identify specific metabolic processes pointing to underlying mechanisms distinguishing them and provide targets for more tailored intervention strategies.

## Materials and methods

The described analysis was performed using *R* version 4.2.1 unless stated otherwise (21).

### Validation RNA-sequencing data processing

RNA-seq raw-counts data from the Genotype-Tissue Expression (GTEx) project (version 8) were downloaded from the *GTEx Portal* on the 25^th^ of November 2022. Samples from subcutaneous adipose tissue, transverse colon, cardiac left ventricle, renal cortex, liver, and skeletal muscle (gastrocnemius) were kept for further analysis. The raw counts data was normalized using geTMM normalization (22), followed by batch effect correction using the *limma* R package (23). As the metabolic task analysis method used, *CellFie* (16), includes logarithmic transformations internally, we shifted every value by 1 to ensure a minimum log-transformed value of 0.

### DCM RNA-sequencing data processing

The Myocardial Applied Genomics Network (MAGNet) RNA-seq raw-counts data (*gene expression omnibus* accession number GSE141910) were downloaded from the consortium’s *GitHub* repository (https://github.com/mpmorley/MAGNet). We only included subjects with a DCM diagnosis or non-failing heart, as determined by the original authors (4). In addition, patients with a reported height under the usual definition of dwarfism (142 cm), although most likely due to human error in reporting, were excluded. The raw count data for the remaining 324 subjects (Supplementary table 1) were then processed as for the GTEx data.

### Metabolic model and Task analysis

All metabolic task analyses were performed using *Matlab* version 2021b and the *Cobra Toolbox version 3.1* (24,25). We used the Human-GEM genome-scale metabolic model version 1.13.0 modified to fit the *Cobra Toolbox* conventions (18,20). We further modified the model adding reactions representing the activities of the *RyR* sarcoplasmic calcium channels and the sarcoplasmic Ca2+ ATPase (Supplementary table 2). We curated a list of metabolic tasks by combining previously published lists (16,18,19), modified to be used with our model, as well as tasks related to cardiac-specific function. From this task list, essential reactions (ERs) were determined using the algorithm from the *checkMetabolicTasks* function of the *Cobra Toolbox,* with minor modifications (Supplementary table 3). This function checks whether the metabolic model contains the minimal set of reactions to complete each metabolic task, and essential reactions (ERS) are metabolic chokepoints which must absolutely be active for the task to be completed. These ERs and the gene expression data were used in the CellFie function to produce task activity scores and binary task activity scores (16). In short, these scores are calculated from the average expression of enzymes contributing to ERs, weighted based on enzyme promiscuity. Although derived from gene expression data, task activity scores are best viewed as having arbitrary units and should not be directly compared across tasks. Binary scores define tasks as “active” if the activity score is higher than a set threshold. We then conducted three CellFie analyses. First with the GTEx data, second with all samples of the MAGNet data, and finally with only the DCM samples of the MAGNet data (see next section). Tasks which were not passed in the *checkMetabolicTasks* run (no feasible set of reactions to fulfil the task to begin with), or whose ERs were not related to any gene such as in passive transport reactions, were removed prior to further analysis. As inspired by the original CellFie paper (16), mixed task scores were created for each of these runs by multiplying the task scores with the binary task scores.

### Clustering based on task scores

The described analysis was performed using *R* version 4.2.1 (21). Tasks considered active or inactive in all samples were excluded from clustering to help contrast samples. Principal component analysis (PCA) was used to recover all principal components (PCs) for the mixed task activity scores of the GTEx data explaining over 99.9% of variance. We then used hierarchical clustering with the *Ward.D2* linkage method to cluster the samples based on these PCs. The appropriate number of clusters was determined via bootstrapping using the bootstrapping algorithm implemented in the *fpc* package version 2.2-11 (26). In short, the algorithm performs an initial hierarchical clustering with all the samples for different numbers of clusters k from 2 to 15. Then, 10000 bootstraps are performed in which subjects are randomly sampled and re-clustered. For each value of k, the Jaccard Similarity index between clusters in the original data and closest clusters in the resampled data is calculated (see *fpc* package documentation for further details). We considered clusters to be stable if they obtained a mean Jaccard similarity above 0.6 across bootstraps. Clusters obtained were then assessed visually using a heatmap and numerically by calculating their “purity”, i.e. the percentage of samples belonging to the most common tissue type in the cluster.

Based on the previous results, we applied the same clustering pipeline to the metabolic task activity of DCM samples in the MAGNet dataset. For this analysis, we only included DCM samples when calculating CellFie continuous and binary activity scores. This was done to avoid situations such as a task being considered inactive in all DCM samples due to lower activity scores in all DCM subjects relative to controls, which would affect gene activity thresholds calculated internally. Such tasks may still show significant differences between DCM subjects and may therefore be relevant for identifying subtypes.

### Hypothesis testing

We performed Mann-whitney U-tests between tissue types for the GTEx-derived task activity scores of a selected set of metabolic tasks. We selected these tasks (“aerobic rephosphorylation of ATP from glucose”, “Glycerol-3-phosphate synthesis”, “Fructose to glucose conversion”) due to having predictable differences between tissue types based on previous literature and databases (1,27–29).

We used the Mann-whitney U-test for every task to determine whether the task activity scores were significantly different between controls and DCM subjects, between controls and subjects from each cluster, and between clusters. For the latter, we used the task activity calculated from DCM subjects only to match the clustering methodology. Results are provided for every task although we focused our interpretation on selected tasks of interest. These tasks were selected as being readily interpretable from the task’s essential reactions and being relevant to cardiac energy metabolism. The chosen tasks cover pathways related to energy generation (oxidative phosphorylation, aerobic respiration from glucose/fatty acids, anaerobic rephosphorylation of ATP from glucose, etc), amino acid metabolism, and cardiac-specific functions (calcium handling, myosin contraction) (Supplementary table 4). In addition, tasks which were considered “inactive” were not included, as their score would have derived from genes expressed at a level we would mostly consider as noise. We employed non-parametric statistical tests as we found task activity to typically violate the normality assumption of t-tests (Supplementary figure 1). In addition, we compared the metabotypes in terms of the limited personal and clinical information available to us (age, sex, BMI, ethnicity, diabetes, hypertension, and left ventricular ejection fraction) using Pearson’s Chi-squared tests and Mann-whitney U-test, as appropriate.

### Immune cell-type deconvolution

We applied the *CIBERSORT* pipeline (R package version 0.1.0) (30), combined with their publicly available immune cells signature matrix, on the MaGNet dataset. For this purpose, the data was normalized into fragments per kilobase of transcript per million (FPKM). We compared abundance percentages using between controls and metabotypes as well as between metabotypes using the mann-whitney u-test, corrected for multiple-testing with the Benjamini–Hochberg procedure.

### Differential gene expression analysis

Differential gene expression analysis was performed using the *edgeR* R package version 4.0.3 (31). The gene expression data was filtered to remove lowly expressed genes using the default parameters of the *filterByExpr* function. After calculating trimmed mean of M values (TMM) normalization factors a quasi-likelihood negative binomial generalized log-linear model was fitted to the data. The model used included the experimental batches and the group variable describing subjects as control or members of a DCM cluster. We then performed differential gene expression analysis using the *glmTreat* function, contrasting between control subjects and subjects from each cluster, as well as between cluster subjects. For more stringent results a minimal log2 fold change of log2(1.5) was required. We used the *clusterProfiler* package version 4.10.0 to perform gene set enrichment analysis using gene ontologies for the tested contrasts (32–34).

### Data and software availability

The transcriptomic data and related metadata from GTEx and MaGNET can be obtained from their portal (https://gtexportal.org/home/downloads/adult-gtex/bulk_tissue_expression) and Github repositories (https://github.com/mpmorley/MAGNet), respectively. The scripts and datasets generated and/or analyzed during the current study are available from the corresponding author upon reasonable request.

## Results

### Metabolic task analysis captures metabolic differences between tissues

We first tested the pipeline’s ability to pick up differences between metabolically distinct tissues (subcutaneous adipose, transverse colon, cardiac left ventricle, skeletal muscle, and liver) by clustering the GTEx samples based on task activity. In doing so we produced 7 tissue specific clusters, although some tissues were split over multiple clusters (average purity of 96%; Supplementary figure 2). As expected, we observed that the energy-demanding cardiac and skeletal muscle tissues presented with significantly higher activity for the “aerobic rephosphorylation of ATP from glucose” task (Figure 1, left) (1,27,28). Adipose tissue samples had the highest task activity for the “Glycerol-3-phosphate synthesis” task (Figure 1, middle) (27). Finally, liver tissue samples were the only ones where “Fructose to glucose conversion” appeared active (Figure 1, right), as expected from mRNA-derived values (29). Colon tissue showed consistently low activity scores for these tasks relative to other tissues.

**Figure 1:**
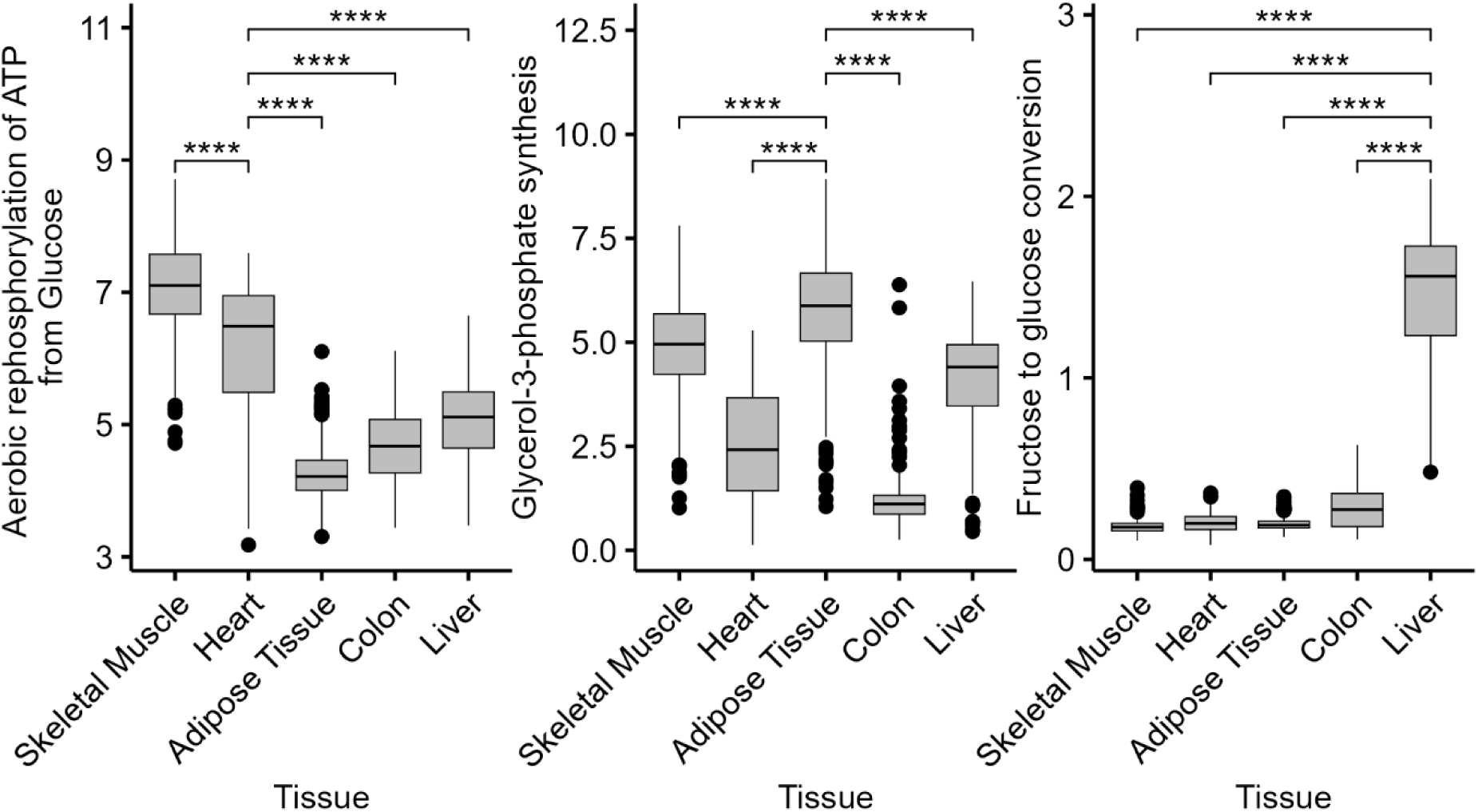
Boxplots of task activity scores (arbitrary units) for three representative tasks, grouped by tissue in the GTEx data. **** p-value < 0.0001.

### End-stage DCM patients separate into two distinct metabotypes

Next, we applied our metabolic tasks analysis pipeline to a publicly available RNA-sequencing dataset of left-ventricular biopsies for 164 DCM patients and 160 non-failing controls (Supplementary table 1). To assess whether subgroups exist within the DCM subjects, we applied our pipeline on the DCM samples separately and identified two stable clusters, from here on referred to as metabotype 1 and metabotype 2 (figure 2; Jaccard coefficients of 0.67 and 0.64 respectively, relative to their closest cluster in each bootstrapped clustering run).

**Figure 2:**
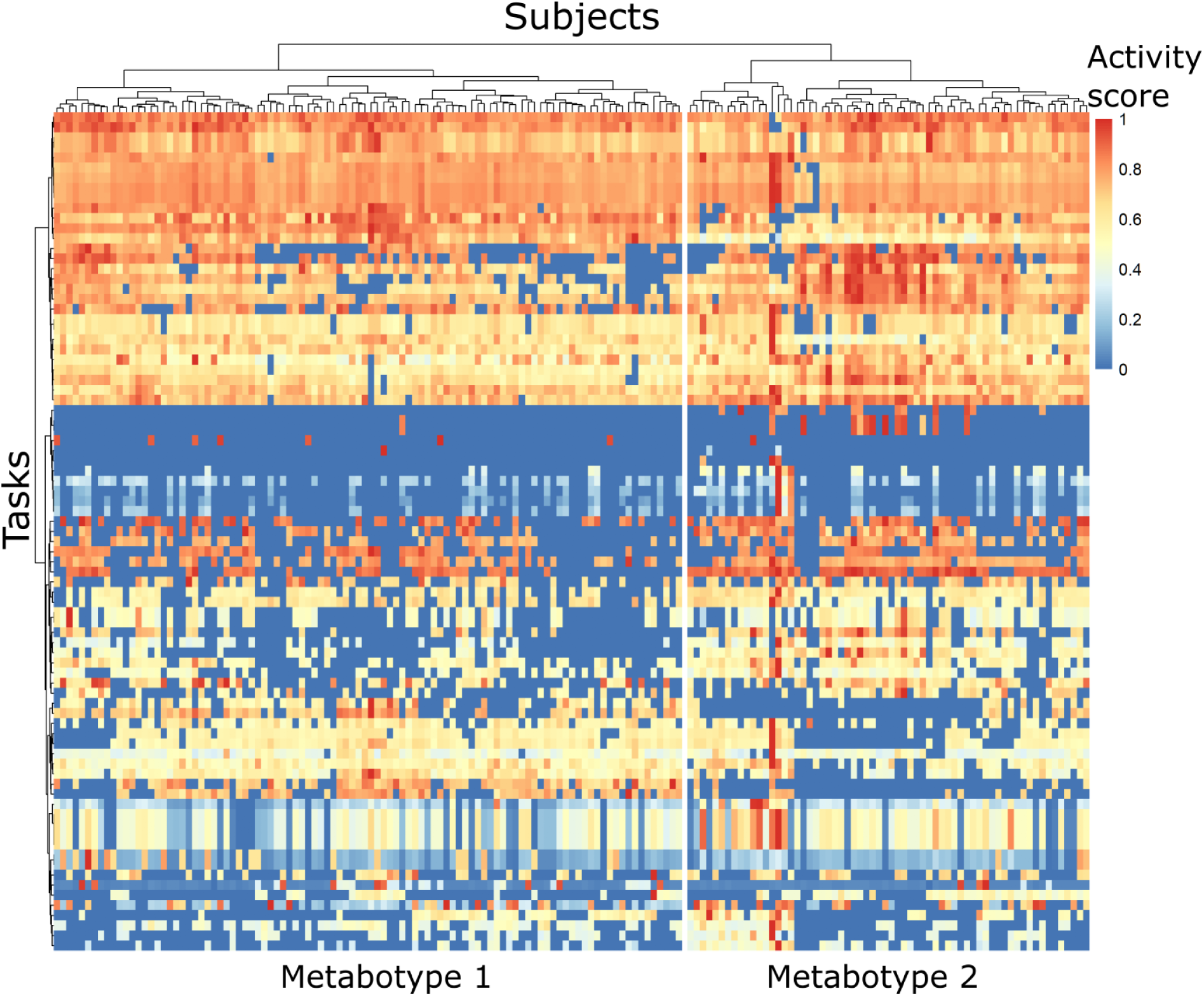
Heatmap of scaled (min-max scaling) mixed task scores of DCM subjects for the tasks included in clustering.

The generated metabotypes presented with distinct metabolic phenotypes based on metabolic task analysis. After accounting for multiple testing, 48 tasks had significantly different activity between metabotypes with a false-discovery rate under 0.05 and absolute fold-change of 10% (Supplementary table 5). Comparisons were also made between control subjects and each metabotype with 43 tasks being significantly altered in metabotype 1 and 54 tasks altered in metabotype 2 (Supplementary table 5). The top 10 significantly different tasks were largely similar for all comparisons involving control subjects, with the task “Carnosine synthesis from beta-alanine and histidine“ being most upregulated in all DCM metabotypes (Supplementary figure 3). By focusing our analysis on predefined tasks of interest, we were able to uncover the distinguishing features that define the metabotypes (Table 1, supplementary figure 4).

**Table 1:**
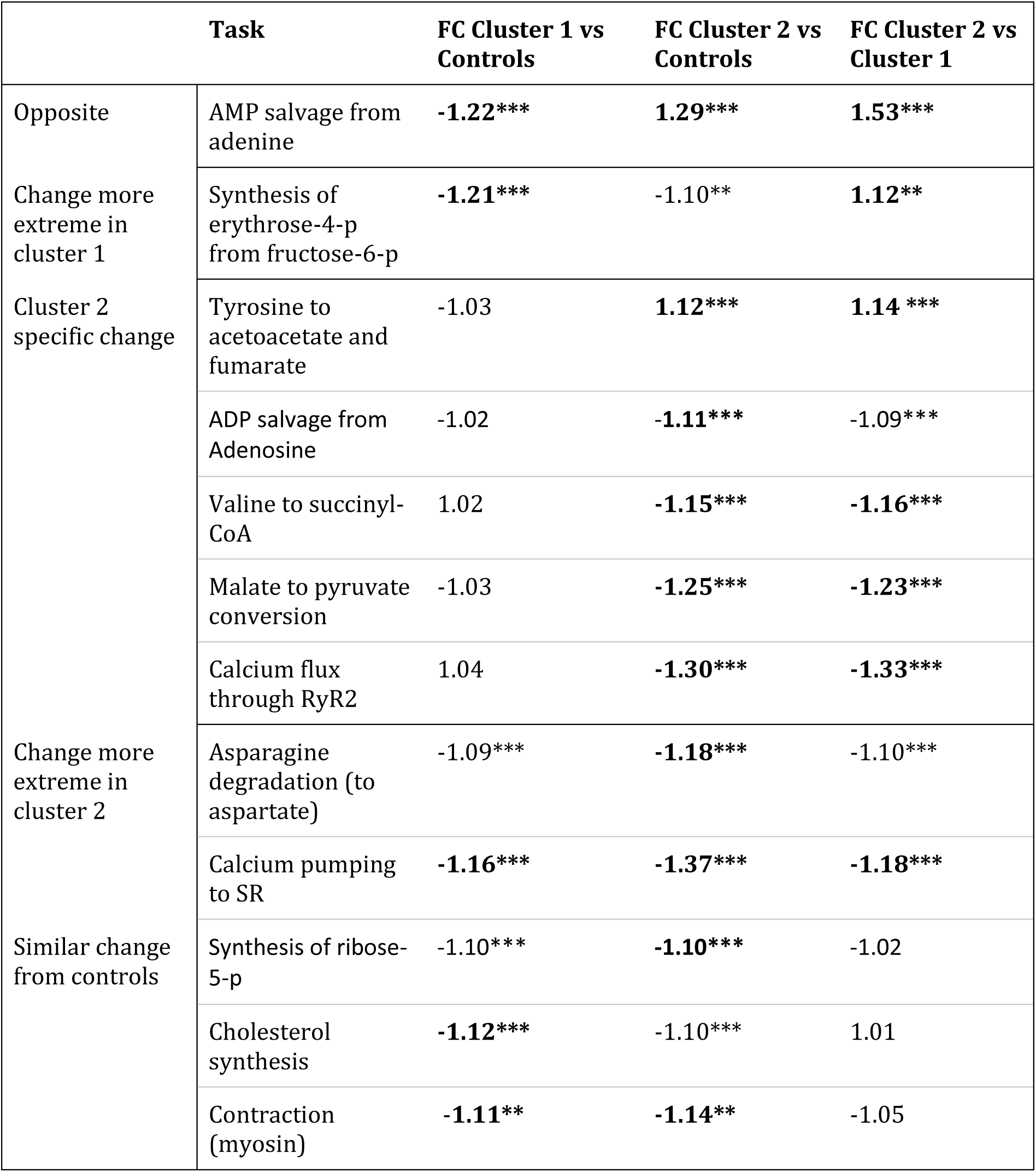
Fold changes of tasks of interest significant in at least one comparison between controls and clusters. * q-value < 0.05, ** q-value < 0.01, *** q-value < 0.001. Bold highlights absolute fold change above 1.1 or below -1.1 (10% difference) in addition to significance. SR = Sarcoplasmic reticulum.

Cardiac-specific tasks were significantly affected in both metabotypes, with downregulation of myosin contraction and calcium pumping into the sarcoplasmic reticulum (SR). Notably, the later was further suppressed in metabotype 2, along with a lower activity of the “Calcium flux through RyR2” task. Tasks related to the pentose phosphate pathway, ribose-5-phosphate and erythrose-4-phosphate synthesis, were downregulated in all DCM subjects, the latter especially in metabotype 1. Amino acid metabolism was especially affected in metabotype 2, with higher activity of “Tyrosine to acetoacetate and fumarate” and lower activity of “Valine to succinyl-CoA”. Asparagine degradation was inhibited in both groups but especially metabotype 2. AMP salvage was downregulated in metabotype 1 and upregulated in metabotype 2, where ADP salvage was lower than in controls. Finally, “Malate to pyruvate conversion” was uniquely downregulated in metabotype 2 while cholesterol synthesis was inhibited in both metabotypes (Table 1).

We did not observe significant differences between metabotypes in terms of the limited available phenotypic and clinical information, including age, sex, ethnicity, BMI, diabetes and hypertension diagnosis, or left-ventricular ejection fraction (LVEF). Median LVEF was 15% in both metabotypes, consistent with the end-stage characterization of these subjects 5 (Table 2).

**Table 2:** Summary characteristics of DCM subjects by metabotype. ^1^n (%); Median (IQR). ^2^Pearson’s Chi-squared test; Wilcoxon rank sum test.

| Characteristic | Metabotype_1, N = 100<sup>1</sup> | Metabotype_2, N = 64<sup>1</sup> | p-value<sup>2</sup> |
| --- | --- | --- | --- |
| Sex: Female | 41 (41%) | 23 (36%) | 0.5 |
| Male | 59 (59%) | 41 (64%) |  |
| Age (years) | 54 (46, 58) | 56 (48, 62) | 0.11 |
| Ethnicity: Afr. American | 46 (46%) | 29 (45%) | >0.9 |
| Caucasian | 54 (54%) | 35 (55%) |  |
| BMI (kg/m<sup>2</sup>) | 25.0 (22.6, 29.3) | 26.7 (24.3, 29.9) | 0.11 |
| Diabetes | 21 (21%) | 18 (28%) | 0.3 |
| Hypertension | 45 (45%) | 32 (50%) | 0.5 |
| LVEF (%): Known | 15 (10, 20) | 15 (10, 20) | 0.5 |
| Unknown | 3 | 2 |  |

### Differential expression analysis suggests a relationship between cardiac metabotypes and the immune system

Next, we extended the analysis of the metabotypes to the whole transcriptome scale by including differential expression analysis to assess differences beyond metabolism. We found 1336 and 2968 differentially expressed genes when comparing controls to metabotype 1 and metabotype 2 respectively (adjusted p < 0.05 and absolute FC > 1.5) (Supplementary figure 5). Comparing controls to metabotypes revealed many enriched lymphocyte-related gene ontologies in metabotype 2, while metabotype 1 presents more enriched ontologies relating to cell-cycle regulation and chemokine production (Figure 3). 1317 genes were differentially expressed between metabotypes, with mostly immune-related ontologies being enriched (Supplementary figure 6).

**Figure 3:**
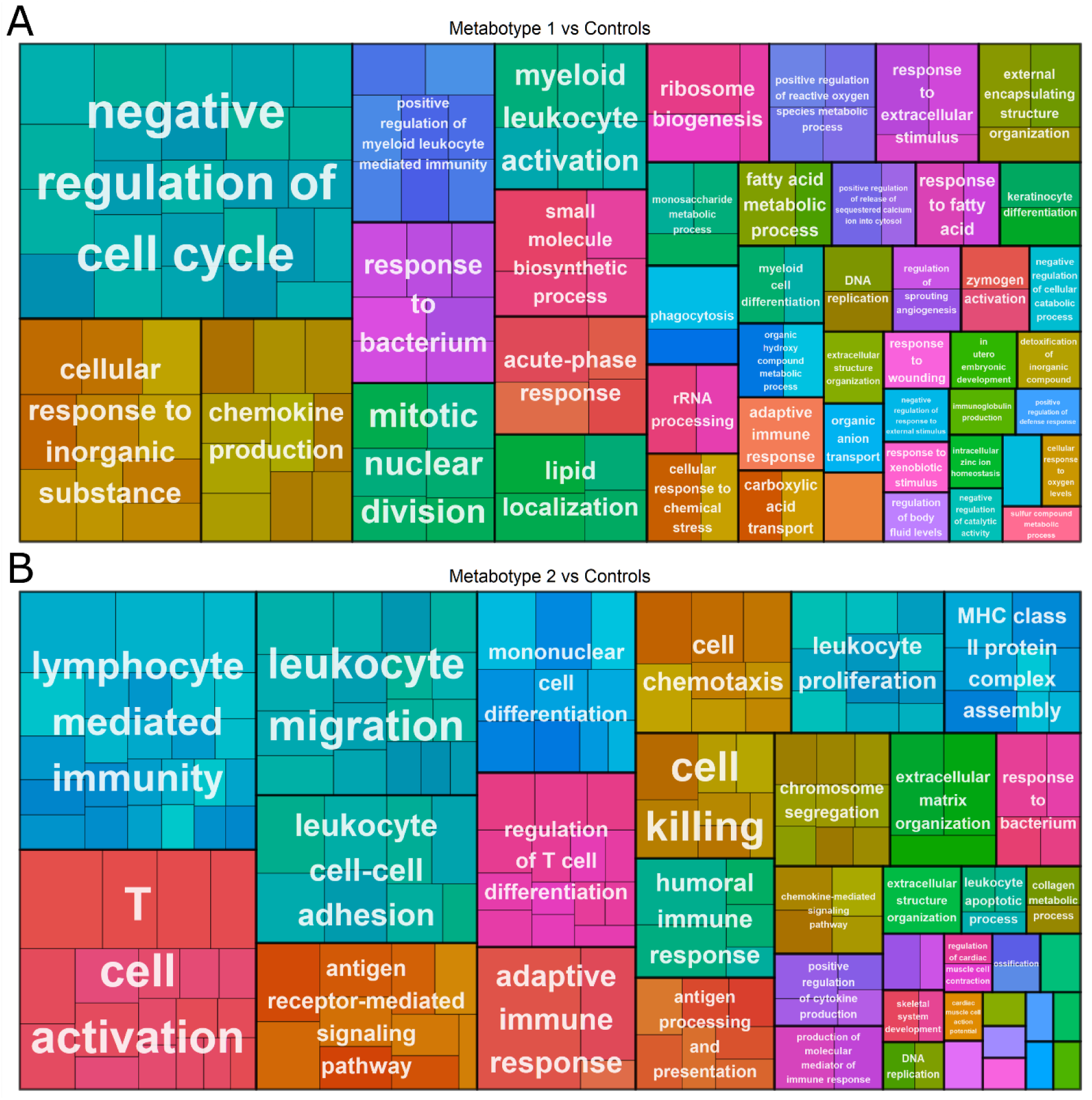
Treemap plot of significantly enriched gene ontologies between controls and metabotypes 1 (A) and 2 (B), grouped by similarity and represented by the most significant ontology in each group. Small rectangles represent individual ontologies with area relative to GSEA score. Colors assigned at random for visual contrast.

### Cell type deconvolution suggests differences in immune cell infiltration between metabotypes

To follow-up on these results, we imputed immune cell-type abundance in DCM samples using the *CIBERSORT* algorithm. Estimated abundance was significantly different between groups for multiple cell-types (Supplementary figure 7). DCM subjects, independently of metabotype, showed an increased proportion of γδ and CD8 T-cells, activated CD4 memory T-cells, activated dendritic cells and B-cells. Further, there were decreases in resting CD4 memory T-cells and memory B-cells. M2 macrophages were lower in DCM subjects as well, and even further in metabotype 1. This metabotype was also characterized by higher proportion of resting dendritic cells as well as lower monocytes. In metabotype 2 we found higher abundance of activated natural killer (NK) cells and lower neutrophils. These results imply immune infiltration differences in DCM patients, most of them consistent between metabotypes.

## Discussion

### Summary of main findings

We constructed a metabotyping pipeline combining established methods and models (16,18,20) which we refined to better suit cardiometabolic research by covering functionalities such as sarcoplasmic calcium handling. We demonstrated the effectiveness of our pipeline using GTEx reference RNA-seq data from different tissues. Accordingly, we showed that our updated pipeline is able to differentiate between tissues both through clustering and through hypothesis testing on tasks related to tissue-specific metabolic requirements. After having validated our pipeline, we applied it to publicly available data of end-stage DCM subjects and control non-failing donors. By doing so, we uncovered two metabotypes in patients with end-stage DCM.

Notably, multiple tasks showed similar directional differences from controls for both metabotypes, even when the activity score was significantly different between metabotypes. Metabotype 1 is characterized by a downregulation of AMP salvage, coupled with a stronger downregulation of the pentose-phosphate pathway (PPP) than in metabotype 2. Metabotype 2, on the other hand, is characterized by changes in a wider range of tasks covering amino acid metabolism, portions of the Krebs cycle and calcium handling-related tasks. Metabotype 2 may therefore represent patients in whom cardiac metabolism is more severely affected than in metabotype 1.

In addition to the observed effects on metabolic functionalities, differential gene expression analysis revealed differences in immune-related pathways: metabotype 1 is associated with cell-cycle inhibition and innate immune responses, while metabotype 2 shows more significant T-cell related pathways. These findings suggest that metabotype 2 may be more affected by immune-related dysregulation, potentially contributing to a more severe disease phenotype.

### Interpretation of main findings

Widespread differential expression in the cardiac transcriptomic profile was observed between controls and end-stage DCM subjects. Dividing the subjects into metabotypes helps to define the heterogeneity in metabolic phenotype displayed in the complete group of DCM patients. Both metabotypes exhibited reduced task activity for the “Contraction” and “Cholesterol synthesis” tasks. Previous research has shown that myosin light-chain kinase may be reduced in end-stage heart failure patients and in left-ventricular assist device non-responders (35,36), while reduced plasma cholesterol is associated with end-stage heart failure, although it is unclear how that would reflect in cardiac cholesterol synthesis (37). Similarly, “ribose-5-phosphate synthesis”, a step of the pentose phosphate pathway (PPP), is downregulated in both metabotypes. The “Synthesis of erythrose-4-p from fructose-6-p” task, which is an entry point into the PPP, is inhibited in both metabotypes, although to a greater extend in metabotype 1. PPP downregulation is in accordance with the observed reduction in intermediate metabolites of this pathway in the subset of these patients analyzed in Flam et al. and may indicate glucose being redirected towards other pathways such as glycolysis (4).

Other tasks, such as “Calcium pumping to SR”, displayed a stronger reduction in metabotype 2 than in metabotype 1. In line with this observation, downregulation of SERCA2, the protein responsible for the active transport of Ca2+ into the sarcoplasmic reticulum in the heart, has been observed both at the mRNA and protein level in ischemic heart failure and idiopathic DCM (38,39). Such a reduction in Ca2+ pumping may limit the Ca2+ concentration in the SR, thus inhibiting contractility through reduced Ca2+ release (39). This may potentially be to a greater extent in metabotype 2 considering the difference in task activity. In addition, the “calcium flux through RyR2” task is significantly reduced in metabotype 2. RyR2 protein expression has been observed in HF (40). However, post-transcriptional modifications of RyR2, which we do not address in this study, have been much more extensively related to HF by impairing calcium handling (40). We suspect that parallel dysregulation of calcium pumping into and release from the SR may contribute to calcium handling impairments, which may contribute to the HF pathology in metabotype 2.

Metabotype 2, compared to controls, shows changes in amino acid metabolism, including a decrease in asparagine and valine degradation and an increase in tyrosine degradation. Lower aspartate levels, a product of asparagine degradation, are linked to poorer heart failure prognosis (41). Branched-chain amino acids (BCAAs) like valine play a key role in heart function, with impaired valine oxidation possibly predicting major adverse cardiac events in non-ischemic DCM (42–44). This study is the first, to our knowledge, to connect tyrosine metabolism to DCM, suggesting an increased demand for ketone bodies as a substrate, supported by previous mouse model research (45).

Finally, the salvage of AMP from adenine was significantly different to controls in opposing directions in both clusters, being increased in metabotype 2 but reduced in metabotype 1. In addition, salvage of ADP from adenosine (with AMP as intermediary product) was found down regulated in metabotype 2. Our results for metabotype 1 can be related to the 2022 paper by Flam and collaborators, which used a subset of the MaGNet dataset used here. They observed in DCM a downregulation of the adenine phosphoribosyltransferase (APRT) enzyme, responsible for the AMP salvage reaction, as well as a decrease of tissue adenine (4). From our literature search, we found no mention of APRT upregulation in the heart, only that it promotes recovery of cutaneous wounds by modulating the ATP/AMP ratio (52). This may imply a link between APRT upregulation and cardiac tissue repair in metabotype 2, though more evidence is needed to support this hypothesis. It is also possible that this result reflects a compensatory effect in metabotype 2 instead.

Both AMP and ADP salvage relate to the purine salvage pathway which is crucial to energy homeostasis in cells as it is partially responsible for maintaining proper ATP/AMP ratio. Dysregulation of this ratio can have broad consequences on the balance of anabolic and catabolic metabolism through AMP-activated protein kinase (AMPK) (46). AMPK-deficient mice models have been shown to develop DCM symptoms (47). Similarly, AMPK inhibition is postulated as a mechanism in doxorubicin-induced DCM (48). Conversely, AMPK activation is regarded as a beneficial therapeutic strategy in cardiac hypertrophy and ischemic heart disease (49,50). However, to our knowledge, the role of AMPK in idiopathic DCM is not quite established yet.

Taken together, results in tasks of interest imply that metabotype 2 differs more from controls relative to metabotype 1, which may indicate a more severe HF. However, it is important to stress that all DCM subjects were already classified as end-stage, requiring cardiac transplant, and that LVEF was similar in both metabotypes. The hypothesized advanced metabolic phenotype of metabotype 2 is therefore more likely to reflect a difference in underlying etiology rather than in disease severity.

Following the metabolic task analysis, we performed usual differential expression analysis and gene-set enrichment analysis (GSEA) to investigate whether metabotypes reflected differences in the whole transcriptome. Most of the metabolic particularities of metabotype 2 observed in metabolic task analysis are not visible in the follow up gene-set enrichment analysis performed. We believe that this highlights the usefulness of metabolic task analysis to detect effects which may go unnoticed in traditional pathway or ontology analysis.

The overwhelming majority of differential expression occurs in genes related to the immune system and inflammation. Metabotype 1 is characterized by ontologies related to cell-cycle inhibition and innate immune response. Cell-cycle regulation, specifically from the Myc gene, has been implicated in various cardiomyopathy animal models and in hypertrophic cardiomyopathy where it is believed to be part of a pathologic “fetal re-expression” pattern (51). Metabotype 2, on the other hand, is heavily characterized by T-cell related ontologies.

As it has been reported before, 9 to 50% of DCM cases involve cardiac inflammation, or myocarditis (52). We know that a wide range of immune cells are involved in inflammation in DCM with what is believed to be a progression from chronic inflammation to cardiac remodeling (53,54). Some studies, however, highlight T-cells as having a particular importance in DCM pathophysiology. Indeed, detection of T-cells in the myocardium is an important factor in diagnosing myocarditis, both in the clinic and in scientific research (55–57). Further, T-cells are directly involved in the pathophysiology of inflammatory DCM. In particular, Th17 CD4+ T-cells promote myocarditis progression into DCM through IL-17A production (58,59). In addition, heart-specific CD4+ T-cells, activated through cross-reactivity to microbiota-derived peptides, can trigger lethal inflammatory cardiomyopathy in mice models and are proposed as a cause for inflammatory cardiomyopathies (60). The observed T-cell-related transcriptomic changes in metabotype 2 may reflect such mechanisms being at play in this patient subgroup.

To investigate this further, we performed cell-type deconvolution analysis on the DCM and control data. The results suggests differences in immune cell infiltration between metabotypes. In metabotype 1 estimated immune cells proportions were lower for monocytes and M2 macrophages, while dendritic cells were increased. Metabotype 2 was characterized by higher NK cells and lower neutrophils. While gene expression-based cell-type estimation has inherent limitations, including its relative nature, these findings highlight broad immune shifts between metabotypes. Interestingly, T-cell subtypes were not significantly different between metabotypes. The T-cell-related transcriptomic signature of metabotype 2 may therefore be related to activity rather than infiltration. This appears to be linked, or at least co-occurring, with metabolic effects such as dysregulation of amino acid metabolism and the purine salvage pathways. This immunometabolic angle is worth investigating further in future studies.

### Limitations and future outlooks

In this work, several technical considerations must be addressed. The effectiveness of metabolic task analysis is primarily constrained by the task list employed in its quality and coverage of human metabolism. While defining metabolic tasks is conceptually straightforward, it is fairly labor-intensive. Nevertheless, we believe that well-defined tasks enhance the efficacy of the pipeline significantly. Moreover, once defined, these tasks can be reused across multiple datasets, increasing utility. By improving and expanding existing task lists, such as our own, the presented pipeline could be applied to a broader range of research applications. In its current implementation though, task activity can already be used in machine learning and multi-omics approaches to identify subtypes, build classifiers and establish links between different molecular modalities.

The clinical relevance of our investigation was also limited by the lack of detailed clinical information on the subjects in the used DCM transcriptomics dataset. While metabotypes did not significantly differ for any clinical variable including left-ventricular ejection fraction, they may still overlap or interact with other information that would be available to clinicians. Our results indicate a distinction in disease severity based on metabolic profiling (e.g., oxidative phosphorylation, amino acid metabolism, and calcium pumping to SR). If metabotypes would be complementary to the clinical information in determining the disease stage, this would have important clinical value in risk prediction of patients with DCM. Follow-up avenues such as a connection between metabotypes and insulin sensitivity or patient outcome cannot be examined without access to more data than was made publicly available. While our results cannot therefore be readily translated to clinical considerations yet, they demonstrate the strength of a metabolic focus in identifying subgroups, confirm existing knowledge and provide novel insights and hypothesis. Future research could focus on experimental confirmation of metabolic task analysis results. For example, one could investigate the increased production of carnosine, a dipeptide with many hypothesized functions in the heart such as antioxidant, pH buffer, and inotropic agent (61), found in all DCM metabotypes and its potential implications. We also advocate for performing this analysis in subjects with varying, known disease severities including early disease stages. This would improve our understanding of cardiometabolic changes before patients progress to end-stage HF. By using data from deeper-phenotyped cohorts we may even be able to find clinical markers reflecting cardiometabolism without the need for endomyocardial biopsies. Further, relating metabolic task activity to inflammatory biomarkers and immunohistological parameters would shine some light onto the immunometabolic differences hinted at by our results. Therapies that target the dysregulated metabolic pathways would be the ultimate goal to improve the outcome for patients with DCM.

## Conclusion

Our approach shows that metabolic task analysis offers an exciting opportunity to uncover insights in DCM, paving the way for a deeper understanding of the progression and heterogeneity. Our results imply the presence of distinct metabotypes in end-stage DCM. Identifying the cause, impact, and progression of these metabolic differences may be crucial to the research effort on DCM and may lead to new personalized treatments.

## Author contributions

B.S.C.N: conceptualization, formal analysis, methodology, software, visualization, writing—original draft, writing—review and editing. J.A.J.V: supervision, writing—review and editing. S.B: resources, writing—review and editing. E.T.: software, writing—review and editing. J.J.F.P.L: writing—review and editing. M.N.: writing—review and editing. S.H.: supervision, funding acquisition, writing—review and editing. M.B: conceptualization, data curation, methodology, project administration, supervision, writing—original draft, writing—review and editing. M.E.A: conceptualization, funding acquisition, methodology, project administration, supervision, writing—original draft, writing—review and editing

The authors declare no competing interests.

## Supplementary material

**Supplementary table 1:** Summary characteristics of the controls and DCM subjects in the MaGNet dataset.

| Characteristic | DCM, N = 164<sup>1</sup> | NF, N = 160<sup>1</sup> | p-value<sup>2</sup> |
| --- | --- | --- | --- |
| Age | 54 (47, 59) | 57 (51, 65) | 0.001 |
| Sex |  |  | 0.015 |
| Female | 64 (39%) | 84 (53%) |  |
| Male | 100 (61%) | 76 (48%) |  |
| Ethnicity |  |  | <0.001 |
| Afr. American | 75 (46%) | 42 (26%) |  |
| Caucasian | 89 (54%) | 118 (74%) |  |
| Diabetes | 39 (24%) | 37 (23%) | 0.9 |
| Hypertension | 77 (47%) | 98 (61%) | 0.010 |

<sup>1</sup>Median (IQR); n (%)
<sup>2</sup>Wilcoxon rank sum test; Pearson's Chi-squared test

**Supplementary table 2:**
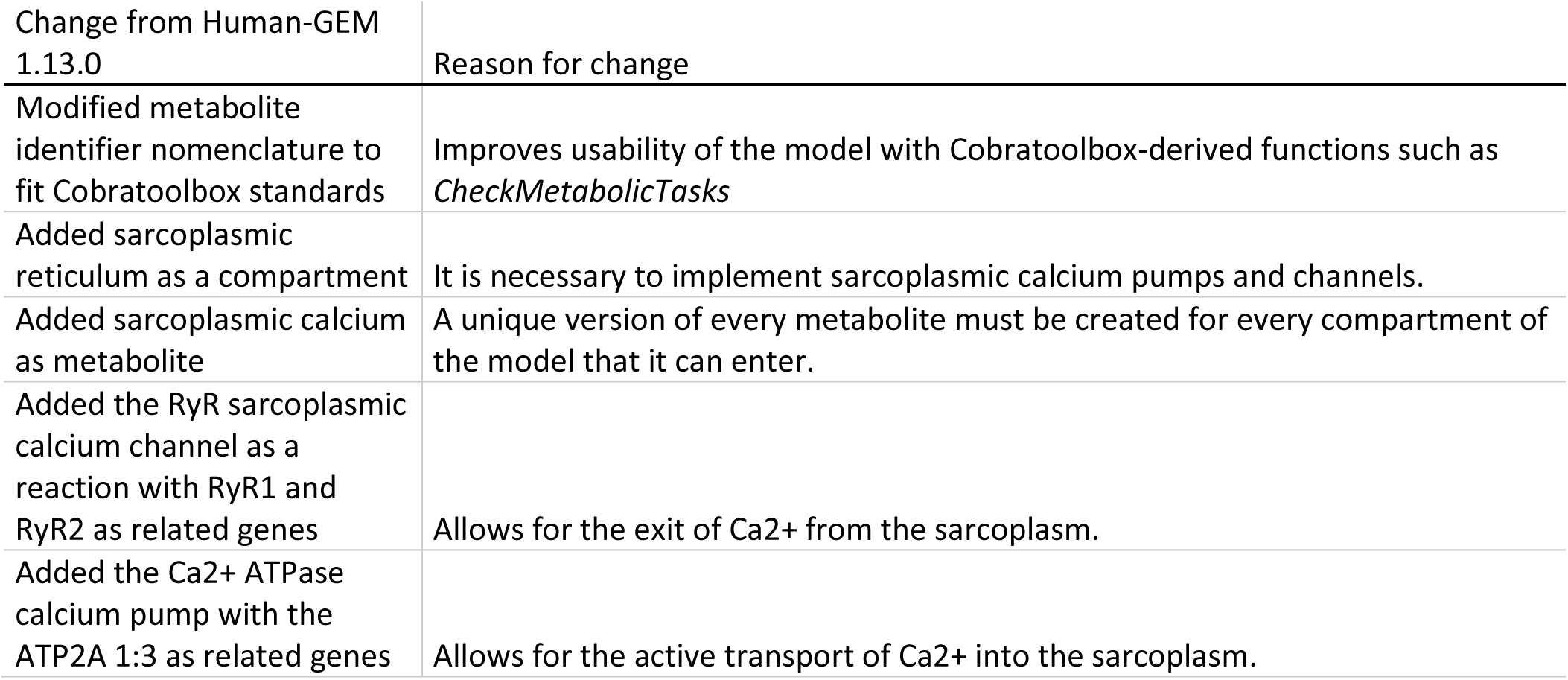
Summary of changes made to the Human-GEM version 1.13.0.

| Change from Human-GEM 1.13.0 | Reason for change |
| --- | --- |
| Modified metabolite identifier nomenclature to fit Cobratoolbox standards | Improves usability of the model with Cobratoolbox-derived functions such as *CheckMetabolicTasks* |
| Added sarcoplasmic reticulum as a compartment | It is necessary to implement sarcoplasmic calcium pumps and channels. |
| Added sarcoplasmic calcium as metabolite | A unique version of every metabolite must be created for every compartment of the model that it can enter. |
| Added the RyR sarcoplasmic calcium channel as a reaction with RyR1 and RyR2 as related genes | Allows for the exit of Ca<sup>2+</sup> from the sarcoplasm. |
| Added the Ca<sup>2+</sup> ATPase calcium pump with the ATP2A 1:3 as related genes | Allows for the active transport of Ca<sup>2+</sup> into the sarcoplasm. |

**Supplementary table 3:**
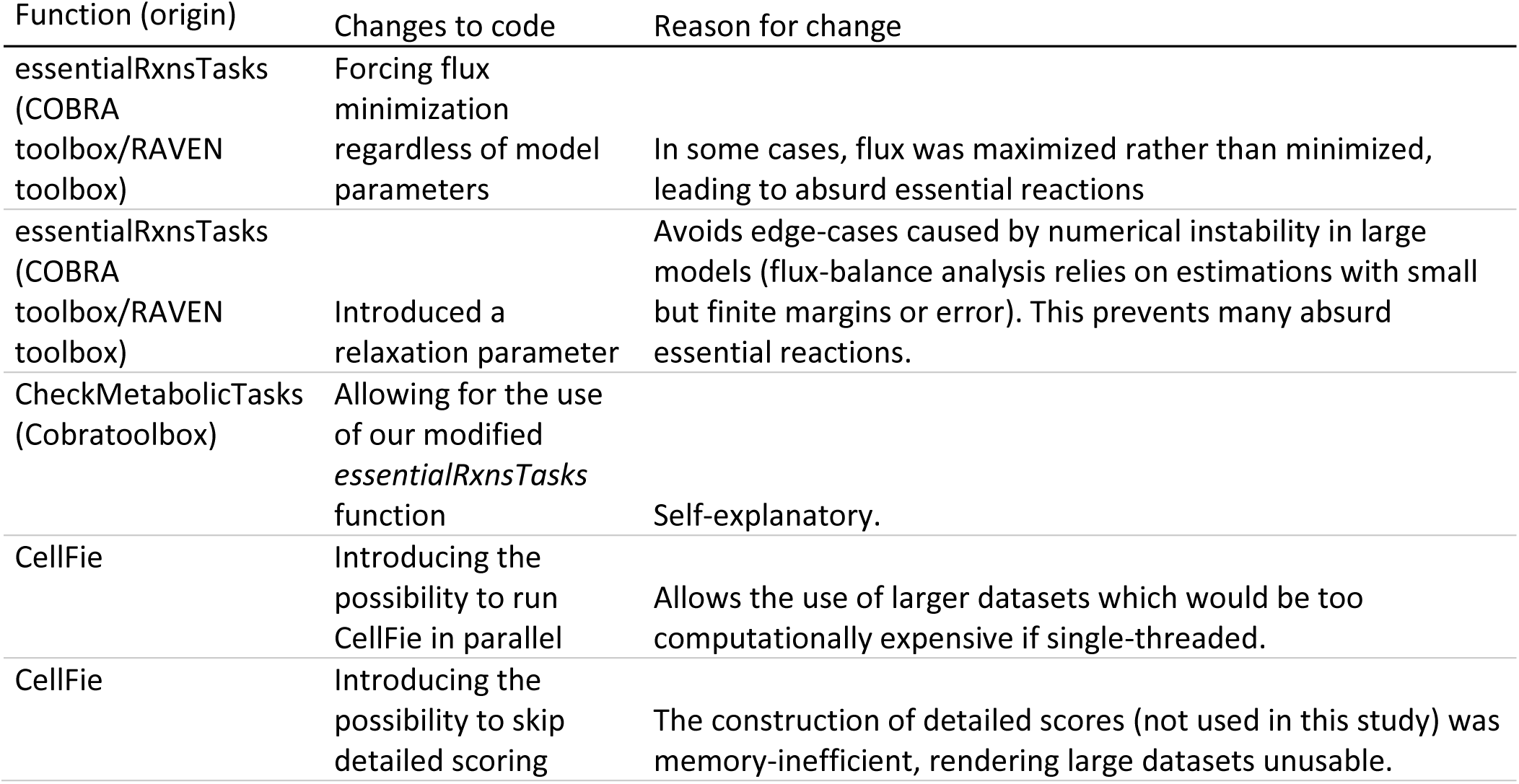
Summary of changes made to Cobratoolbox and Cellfie functions.

| Function (origin) | Changes to code | Reason for change |
| --- | --- | --- |
| essentialRxnsTasks (COBRA toolbox/RAVEN toolbox) | Forcing flux minimization regardless of model parameters | In some cases, flux was maximized rather than minimized, leading to absurd essential reactions |
| essentialRxnsTasks (COBRA toolbox/RAVEN toolbox) | Introduced a relaxation parameter | Avoids edge-cases caused by numerical instability in large models (flux-balance analysis relies on estimations with small but finite margins or error). This prevents many absurd essential reactions. |
| CheckMetabolicTasks (Cobratoolbox) | Allowing for the use of our modified *essentialRxnsTasks* function | Self-explanatory. |
| CellFie | Introducing the possibility to run CellFie in parallel | Allows the use of larger datasets which would be too computationally expensive if single-threaded. |
| CellFie | Introducing the possibility to skip detailed scoring | The construction of detailed scores (not used in this study) was memory-inefficient, rendering large datasets unusable. |

**Supplementary table 4:**
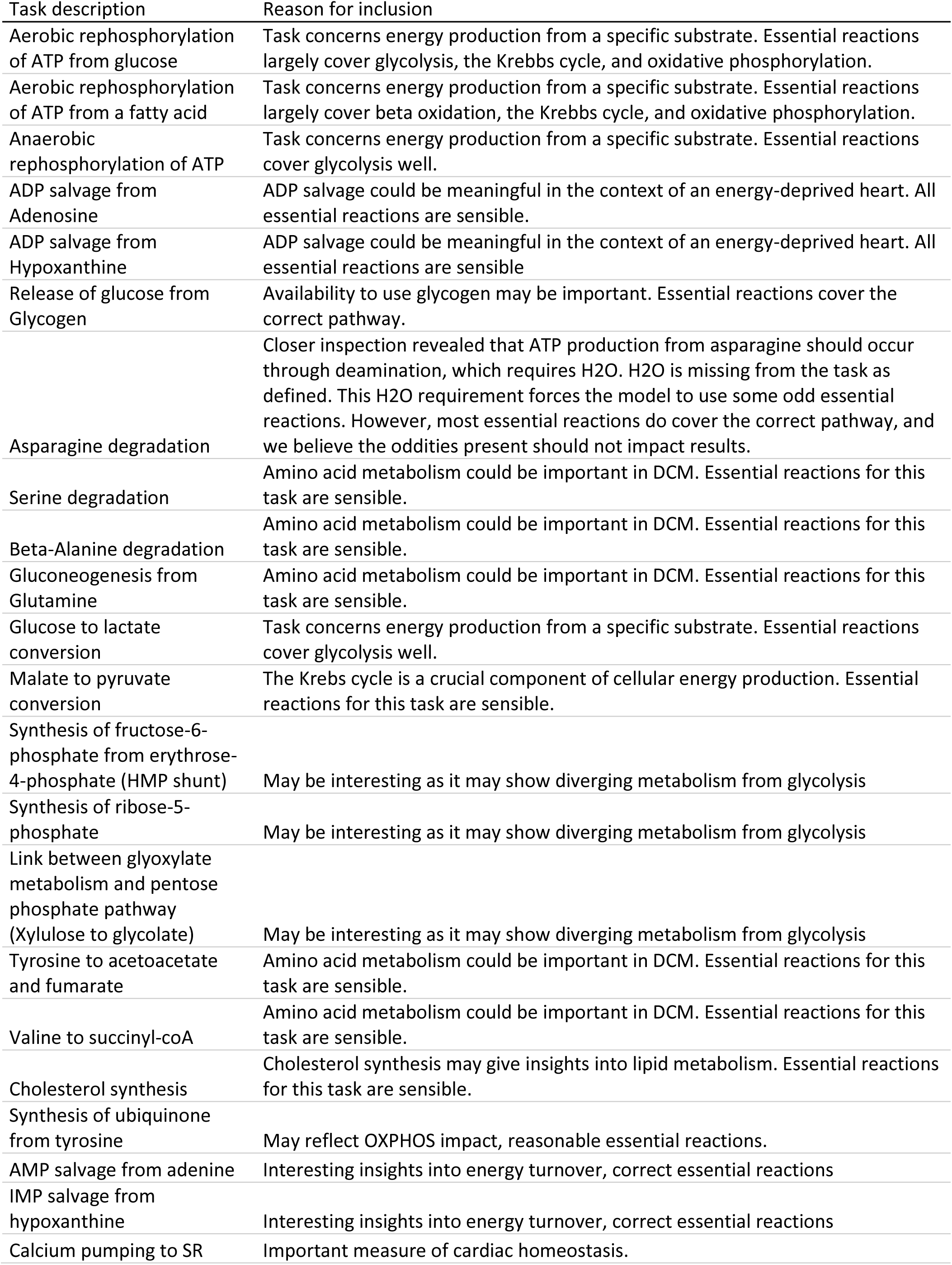
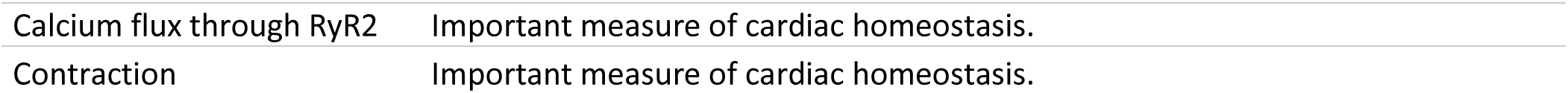
Tasks of interest focused upon in analysis.

**Supplementary figure 1:**
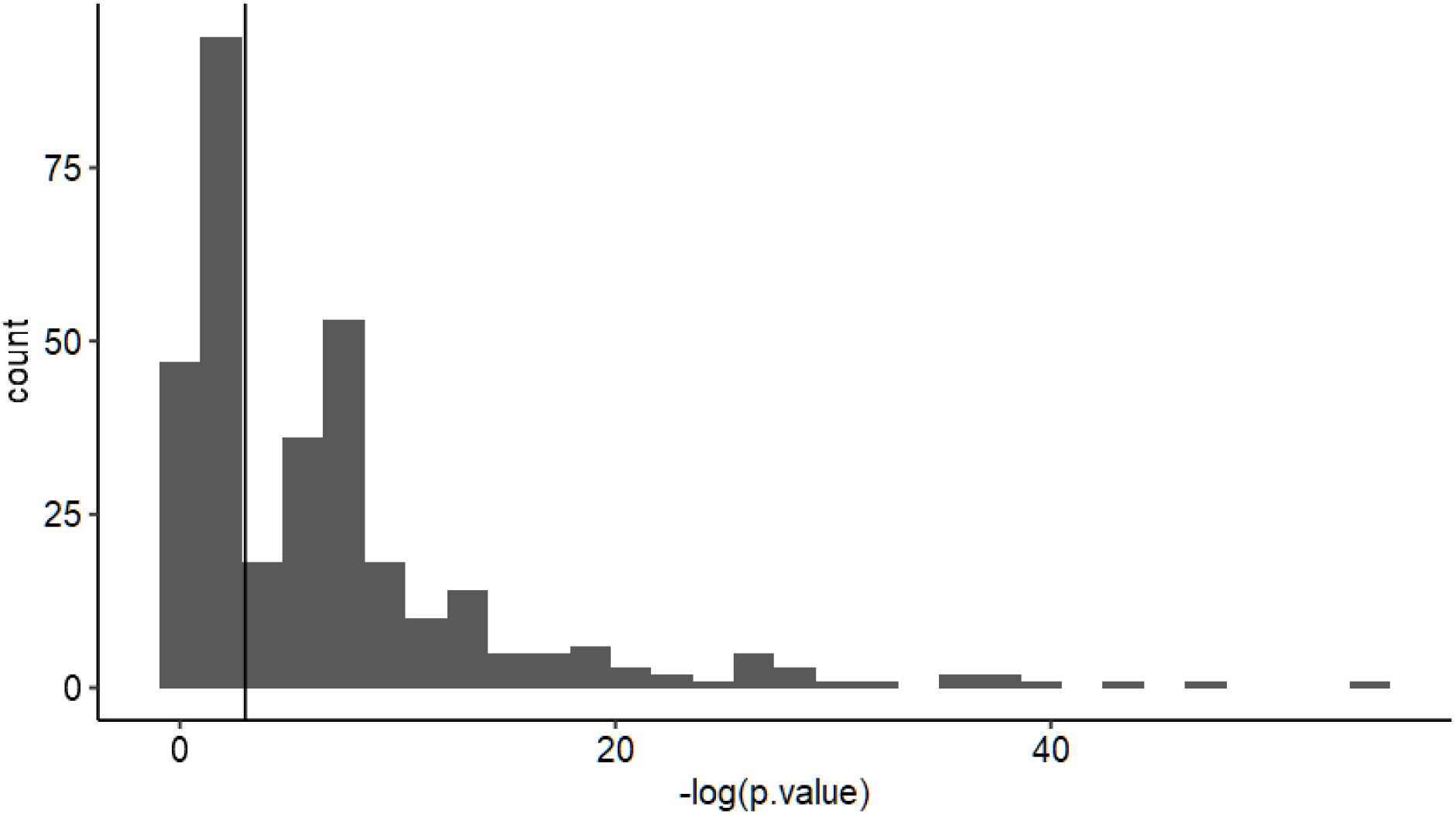
histogram of Shapiro-test p-values, calculated for each task from the task activity score of MaGNet control samples. Values to the left of the vertical line do not fit the normality assumption.

**Supplementary figure 2:**
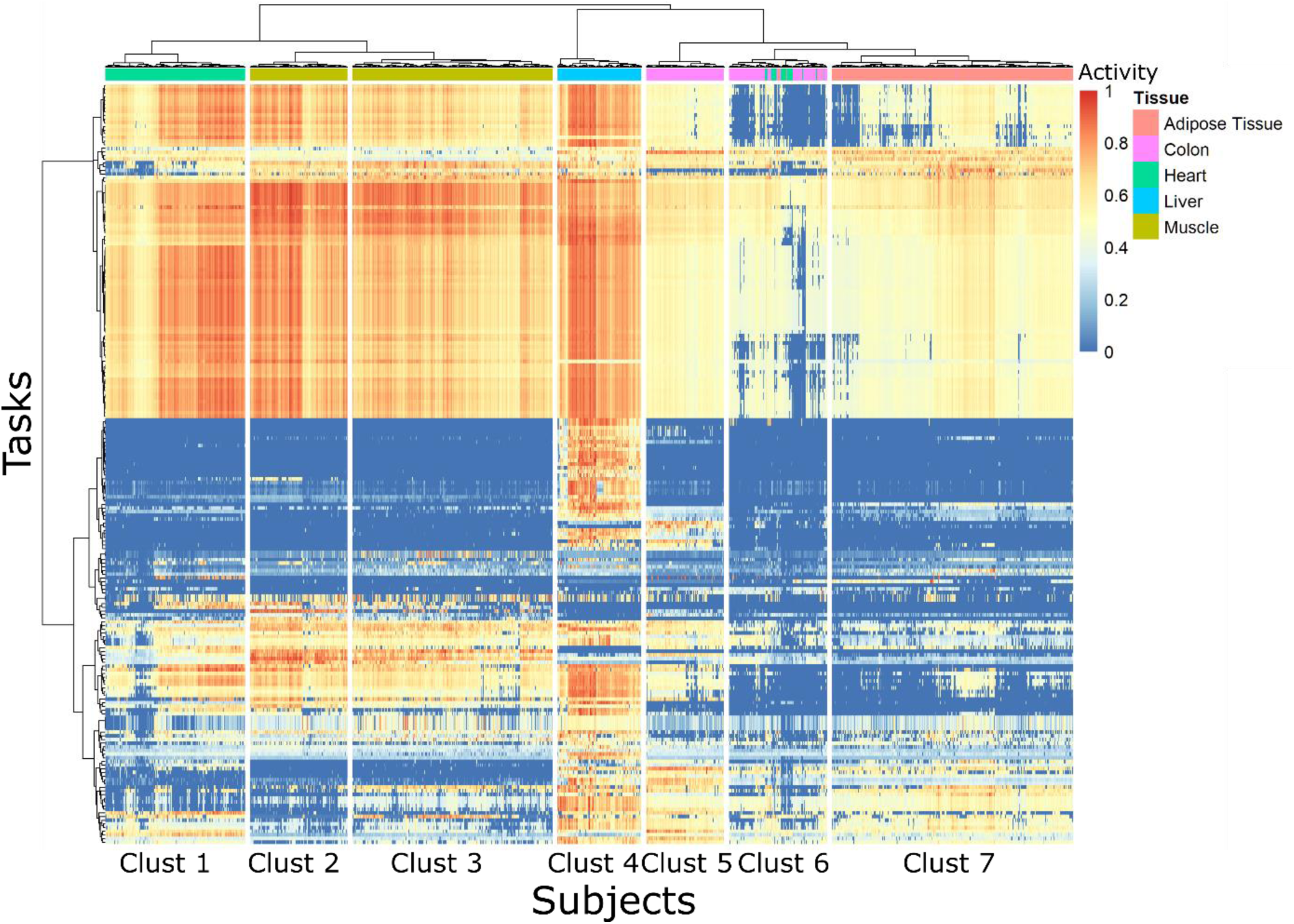
Heatmap of GTEx mixed activity, showing the clustering.

**Supplementary table 5:** Fold changes between controls and clusters for all tasks. * q-value < 0.05, ** q-value < 0.01, *** q-value < 0.001. Bold highlights absolute fold change above 1.1 or below -1.1 (10% difference) in addition to significance. Met. = Metabotype.

| Task | FC DCM vs Controls | FC Met. 1 vs Controls | FC Met. 2 vs Controls | FC Met. 2 vs Met. 1 |
| --- | --- | --- | --- | --- |
| Task_1: Aerobic rephosphorylation of ATP from glucose | -1.01 | -1.02** | 1.01 | 1.02** |
| Task_2: Aerobic rephosphorylation of ATP from a fatty acid | 1.02* | 1.01 | 1.03** | 1.02* |
| Task_3: Aerobic rephosphorylation of GTP | 1.00 | -1.01 | 1.03* | 1.03** |
| Task_4: Aerobic rephosphorylation of CTP | 1.00 | -1.01 | 1.03* | 1.04** |
| Task_5: Aerobic rephosphorylation of UTP | 1.00 | -1.01 | 1.03* | 1.04** |
| Task_6: Anaerobic rephosphorylation of ATP | 1.00 | -1.02 | 1.04** | 1.06*** |
| Task_7: Anaerobic rephosphorylation of GTP | 1.00 | -1.02 | 1.04** | 1.06*** |
| Task_8: Anaerobic rephosphorylation of CTP | 1.00 | -1.02 | 1.04** | 1.06*** |
| Task_9: Anaerobic rephosphorylation of UTP | 1.00 | -1.02 | 1.04** | 1.06*** |
| Task_10: ATP de novo synthesis | -1.03*** | -1.04*** | -1.01 | 1.03** |
| Task_11: CTP de novo synthesis | -1.02* | -1.03* | -1.02 | 1.00 |
| Task_12: GTP de novo synthesis | -1.07*** | -1.07*** | -1.07*** | -1.02* |
| Task_13: UTP de novo synthesis | -1.02 | -1.02 | -1.01 | 1.00 |
| Task_14: dATP de novo synthesis | -1.08*** | -1.07*** | **-1.11\*\*\*** | -1.05*** |
| Task_15: dCTP de novo synthesis | -1.02* | -1.03* | -1.02 | 1.00 |
| Task_16: dGTP de novo synthesis | -1.07*** | -1.07*** | -1.08*** | -1.02* |
| Task_17: dTTP de novo synthesis | -1.01 | -1.03* | 1.01 | 1.03 |
| Task_18: ADP salvage from Adenosine | -1.05** | -1.02 | **-1.11\*\*\*** | -1.09*** |
| Task_19: ADP salvage from Hypoxanthine | 1.04 | 1.04 | 1.04 | 1.00 |
| Task_20: dTTP salvage from Thymine | -1.05 | **-1.18\*\*\*** | **1.13\*\*** | **1.33\*\*\*** |
| Task_21: Aerobic reduction of NAD+ | -1.01 | -1.03** | 1.01 | 1.04*** |
| Task_22: Aerobic reduction of NADP+ | 1.01 | -1.02 | 1.05* | 1.07*** |
| Task_23: Gluconeogenesis from Lactate | -1.01 | -1.05*** | 1.05*** | 1.09*** |
| Task_24: Gluconeogenesis from Glycerol | -1.00 | -1.05*** | 1.06*** | **1.11\*\*\*** |
| Task_25: Gluconeogenesis from Alanine | -1.01 | -1.05*** | 1.05*** | 1.09*** |
| Task_26: Gluconeogenesis from Lactate and optionally fatty acid | 1.00 | -1.01 | 1.03** | 1.03*** |
| Task_27: Gluconeogenesis from Glycerol and optionally fatty acid | -1.01 | -1.03** | 1.02 | 1.04*** |
| Task_28: Gluconeogenesis from Alanine and fatty acid | -1.02* | -1.03*** | 1.00 | 1.03* |
| Task_29: Storage of glucose in Glycogen | NA | NA | NA | NA |
| Task_30: Release of glucose from Glycogen | 1.00 | -1.00 | 1.01 | 1.00 |
| Task_31: Fructose degradation | -1.01 | -1.02** | 1.00 | 1.02** |
| Task_32: Galactose degradation | -1.01 | -1.02** | 1.01 | 1.02** |
| Task_33: UDP-glucose de novo synthesis | -1.09*** | -1.06*** | **-1.14\*\*\*** | -1.08*** |
| Task_34: UDP-galactose de novo synthesis | 1.07* | 1.10** | 1.04 | -1.06 |
| Task_35: UDP-glucuronate de novo synthesis | -1.09*** | -1.08*** | **-1.11\*\*\*** | -1.04** |
| Task_36: GDP-L-fucose de novo synthesis | 1.02 | 1.00 | 1.05 | 1.03 |
| Task_37: GDP-mannose de novo synthesis | 1.03 | 1.04 | 1.03 | -1.04 |
| Task_38: UDP-N-acetyl D-galactosamine de novo synthesis | -1.09*** | -1.10*** | -1.08*** | 1.02 |
| Task_39: CMP-N-acetylneuraminate de novo synthesis | -1.10*** | **-1.11\*\*\*** | -1.09*** | 1.02 |
| Task_40: N-Acetylglucosamine de novo synthesis | -1.06*** | -1.08*** | -1.02 | 1.05** |
| Task_41: Glucuronate de novo synthesis | -1.09*** | -1.07*** | **-1.11\*\*\*** | -1.04** |
| Task_42: Alanine de novo synthesis | -1.03*** | -1.05*** | -1.00 | 1.04*** |
| Task_43: Arginine de novo synthesis | -1.02*** | -1.04*** | -1.00 | 1.03*** |
| Task_44: Asparagine de novo synthesis | -1.04*** | -1.05*** | -1.03* | 1.02* |
| Task_45: Aspartate de novo synthesis | -1.02* | -1.04*** | 1.00 | 1.03** |
| Task_46: Glutamate de novo synthesis | -1.03*** | -1.05*** | -1.01 | 1.03*** |
| Task_47: Glycine de novo synthesis | -1.02** | -1.03*** | -1.00 | 1.03*** |
| Task_48: Glutamine de novo synthesis | -1.05*** | -1.06*** | -1.03* | 1.03** |
| Task_49: Proline de novo synthesis | -1.02 | -1.03*** | 1.01 | 1.03** |
| Task_50: Serine de novo synthesis | -1.01 | -1.02** | 1.01 | 1.02** |
| Task_51: Histidine de novo synthesis | NA | NA | NA | NA |
| Task_52: Histidine uptake | -1.00 | -1.09* | **1.12** | **1.24\*\*** |
| Task_53: Isoleucine de novo synthesis | NA | NA | NA | NA |
| Task_54: Isoleucine uptake | -1.00 | -1.09* | **1.12** | **1.24\*\*** |
| Task_55: Leucine de novo synthesis | NA | NA | NA | NA |
| Task_56: Leucine uptake | -1.00 | -1.09* | **1.12** | **1.24\*\*** |
| Task_57: Lysine de novo synthesis | NA | NA | NA | NA |
| Task_58: Lysine uptake | **-1.18\*\*** | **-1.24\*\*\*** | -1.09 | **1.37\*\*** |
| Task_59: Methionine de novo synthesis | NA | NA | NA | NA |
| Task_60: Methionine uptake | -1.05*** | -1.06** | -1.03* | 1.00 |
| Task_61: Phenylalanine de novo synthesis | NA | NA | NA | NA |
| Task_62: Phenylalanine uptake | **1.27** | **1.36** | **1.14** | **-1.38** |
| Task_63: Threonine de novo synthesis | NA | NA | NA | NA |
| Task_64: Threonine uptake | 1.06 | -1.09* | **1.28** | **1.38\*\*** |
| Task_65: Tryptophan de novo synthesis | NA | NA | NA | NA |
| Task_66: Tryptophan uptake | **1.27** | **1.36** | **1.14** | **-1.38** |
| Task_67: Valine de novo synthesis | NA | NA | NA | NA |
| Task_68: Valine uptake | -1.00 | -1.09* | **1.12** | **1.24\*\*** |
| Task_69: Cysteine de novo synthesis from glucose | NA | NA | NA | NA |
| Task_70: Cysteine de novo synthesis from AAs | **-1.35\*\*\*** | **-1.34\*\*\*** | **-1.37\*\*\*** | -1.02 |
| Task_71: Cystine de novo synthesis | NA | NA | NA | NA |
| Task_72: Tyrosine de novo synthesis | NA | NA | NA | NA |
| Task_73: Tyrosine de novo synthesis | 1.09*** | 1.07*** | **1.12\*\*\*** | 1.04* |
| Task_74: Homocysteine de novo synthesis | NA | NA | NA | NA |
| Task_75: Homocysteine de novo synthesis | **-1.34\*\*\*** | **-1.32\*\*\*** | **-1.38\*\*\*** | -1.03 |
| Task_76: beta-Alanine de novo synthesis | -1.03*** | -1.04*** | -1.02 | 1.01 |
| Task_77: Alanine degradation | -1.07*** | -1.06*** | -1.08*** | -1.04** |
| Task_78: Arginine degradation | -1.04*** | -1.05*** | -1.03** | 1.01 |
| Task_79: Asparagine degradation | **-1.12\*\*\*** | -1.09*** | **-1.17\*\*\*** | **-1.10\*\*\*** |
| Task_80: Aspartate degradation | -1.06*** | -1.07*** | -1.06*** | -1.01 |
| Task_81: Cysteine degradation | -1.06*** | -1.05*** | -1.06*** | -1.02 |
| Task_82: Glutamate degradation | -1.07*** | -1.07*** | -1.06*** | 1.00 |
| Task_83: Glycine degradation | 1.02 | -1.05* | **1.14\*\*\*** | **1.19\*\*\*** |
| Task_84: Histidine degradation | -1.07*** | -1.07*** | -1.06*** | 1.00 |
| Task_85: Isoleucine degradation | -1.06*** | -1.06*** | -1.05*** | -1.00 |
| Task_86: Glutamine degradation | -1.06*** | -1.06*** | -1.05*** | 1.00 |
| Task_87: Leucine degradation | -1.05*** | -1.05*** | -1.06*** | -1.02 |
| Task_88: Lysine degradation | -1.06*** | -1.06*** | -1.05*** | -1.00 |
| Task_89: Methionine degradation | -1.06*** | -1.07*** | -1.04** | 1.02* |
| Task_90: Phenylalanine degradation | -1.06*** | -1.06*** | -1.05*** | 1.00 |
| Task_91: Proline degradation | -1.05*** | -1.06*** | -1.05*** | 1.00 |
| Task_92: Serine degradation | -1.03 | -1.07*** | 1.04* | **1.12\*\*\*** |
| Task_93: Threonine degradation | -1.06*** | -1.06*** | -1.06*** | -1.01 |
| Task_94: Tryptophan degradation | -1.05*** | -1.06*** | -1.04*** | 1.00 |
| Task_95: Tyrosine degradation | -1.06*** | -1.06*** | -1.07*** | -1.03* |
| Task_96: Valine degradation | -1.05*** | -1.06*** | -1.05*** | -1.00 |
| Task_97: Homocysteine degradation | **-1.42\*\*\*** | **-1.47\*\*\*** | **-1.35\*\*\*** | 1.07* |
| Task_98: Ornithine degradation | -1.05*** | -1.06*** | -1.05*** | 1.00 |
| Task_99: beta-Alanine degradation | -1.06*** | -1.05*** | -1.07*** | -1.03 |
| Task_100: Urea from alanine | -1.04*** | -1.04*** | -1.03* | 1.01 |
| Task_101: Urea from glutamine | -1.04*** | -1.04*** | -1.03** | -1.00 |
| Task_102: Creatine de novo synthesis | **1.43\*\*\*** | **1.39\*\*\*** | **1.50\*\*\*** | 1.09 |
| Task_103: Heme de novo synthesis | -1.04*** | -1.02** | -1.06*** | -1.04*** |
| Task_104: PC de novo synthesis | -1.04*** | -1.04*** | -1.03** | 1.01 |
| Task_105: PE de novo synthesis | -1.04*** | -1.05*** | -1.02* | 1.02** |
| Task_106: PS de novo synthesis | -1.03*** | -1.04*** | -1.02* | 1.01 |
| Task_107: PI de novo synthesis | -1.03*** | -1.04*** | -1.01 | 1.02* |
| Task_108: Cardiolipin de novo synthesis | -1.02*** | -1.03*** | -1.01 | 1.01 |
| Task_109: SM de novo synthesis | -1.04*** | -1.04*** | -1.03*** | 1.00 |
| Task_110: Ceramide de novo synthesis | -1.03*** | -1.04*** | -1.02** | 1.01 |
| Task_111: Lactosylceramide de novo synthesis | -1.03*** | -1.03*** | -1.02** | -1.00 |
| Task_112: CoA de novo synthesis | 1.03* | -1.01 | **1.10\*\*\*** | **1.11\*\*\*** |
| Task_113: NAD de novo synthesis | **-1.43\*\*\*** | **-1.31\*\*\*** | **-1.66\*\*\*** | **-1.25\*\*\*** |
| Task_114: NADP de novo synthesis | -1.02** | -1.04*** | -1.00 | 1.03* |
| Task_115: FAD de novo synthesis | **-1.16\*\*\*** | **-1.15\*\*\*** | **-1.17\*\*\*** | -1.02 |
| Task_116: Thioredoxin de novo synthesis | NA | NA | NA | NA |
| Task_117: THF de novo synthesis | -1.08** | -1.03 | **-1.15\*\*\*** | **-1.11\*\*\*** |
| Task_118: Pyridoxal-P de novo synthesis | -1.04* | **-1.12\*\*\*** | 1.06 | **1.17\*\*\*** |
| Task_119: Acetoacetate de novo synthesis | -1.05*** | -1.06*** | -1.03* | 1.02 |
| Task_120: (R)-3-Hydroxybutanoate de novo synthesis | -1.04*** | -1.05*** | -1.03 | 1.02 |
| Task_121: Farnesyl-PP de novo synthesis | -1.01 | -1.08*** | 1.09*** | **1.16\*\*\*** |
| Task_122: Lauric acid de novo synthesis | -1.03*** | -1.03*** | -1.02 | 1.01 |
| Task_123: Tridecylic acid de novo synthesis | -1.03*** | -1.04*** | -1.02 | 1.02* |
| Task_124: Myristic acid de novo synthesis | -1.03*** | -1.03*** | -1.02 | 1.01 |
| Task_125: 9E-tetradecenoic acid de novo synthesis | -1.03*** | -1.03*** | -1.02 | 1.01 |
| Task_126: 7Z-tetradecenoic acid de novo synthesis | -1.03*** | -1.03*** | -1.02 | 1.01 |
| Task_127: Physeteric acid de novo synthesis | -1.03*** | -1.03*** | -1.02 | 1.01 |
| Task_128: Pentadecylic acid de novo synthesis | -1.03*** | -1.04*** | -1.02 | 1.02* |
| Task_129: Palmitate de novo synthesis | -1.03*** | -1.04*** | -1.02 | 1.02* |
| Task_130: Palmitolate de novo synthesis | -1.03** | -1.04*** | -1.01 | 1.02* |
| Task_131: 7-palmitoleic acid de novo synthesis | -1.03*** | -1.03*** | -1.02 | 1.01 |
| Task_132: Margaric acid de novo synthesis | -1.03*** | -1.04*** | -1.02 | 1.02* |
| Task_133: 10Z-heptadecenoic acid de novo synthesis | -1.03*** | -1.04*** | -1.02 | 1.02* |
| Task_134: 9-heptadecylenic acid de novo synthesis | -1.03*** | -1.04*** | -1.02 | 1.02* |
| Task_135: Stearate de novo synthesis | -1.01 | -1.02 | 1.01 | 1.02* |
| Task_136: 13Z-octadecenoic acid de novo synthesis | -1.03*** | -1.03*** | -1.02 | 1.01 |
| Task_137: cis-vaccenic acid de novo synthesis | -1.03** | -1.04*** | -1.01 | 1.02* |
| Task_138: Oleate de novo synthesis | -1.01 | -1.02 | 1.01 | 1.02* |
| Task_139: Elaidate de novo synthesis | -1.03** | -1.04*** | -1.01 | 1.02* |
| Task_140: 7Z-octadecenoic acid de novo synthesis | -1.03*** | -1.03*** | -1.02 | 1.01 |
| Task_141: 6Z,9Z-octadecadienoic acid de novo synthesis | -1.03*** | -1.03*** | -1.02 | 1.01 |
| Task_142: Nonadecylic acid de novo synthesis | -1.03*** | -1.04*** | -1.02 | 1.02* |
| Task_143: Eicosanoate de novo synthesis | -1.03*** | -1.03*** | -1.02 | 1.01 |
| Task_144: 13Z-Eicosenoic acid de novo synthesis | -1.03*** | -1.03*** | -1.02 | 1.01 |
| Task_145: cis-gondoic acid de novo synthesis | -1.03*** | -1.03*** | -1.02 | 1.01 |
| Task_146: 9-Eicosenoic acid de novo synthesis | -1.03*** | -1.03*** | -1.02 | 1.01 |
| Task_147: 8,11-Eicosadienoic acid de novo synthesis | -1.03*** | -1.03*** | -1.02 | 1.01 |
| Task_148: Mead acid de novo synthesis | -1.03*** | -1.04*** | -1.02 | 1.01 |
| Task_149: Henicosanoic acid de novo synthesis | -1.03*** | -1.04*** | -1.02 | 1.02* |
| Task_150: Behenic acid de novo synthesis | -1.03*** | -1.03*** | -1.02 | 1.01 |
| Task_151: cis-erucic acid de novo synthesis | -1.03*** | -1.03*** | -1.02 | 1.01 |
| Task_152: cis-cetoleic acid de novo synthesis | -1.04*** | -1.05*** | -1.04** | 1.01 |
| Task_153: Tricosanoic acid de novo synthesis | -1.03*** | -1.04*** | -1.01 | 1.02* |
| Task_154: Lignocerate de novo synthesis | -1.03*** | -1.03*** | -1.02 | 1.00 |
| Task_155: Nervonic acid de novo synthesis | -1.04*** | -1.05*** | -1.04** | 1.00 |
| Task_156: Cerotic acid de novo synthesis | -1.03*** | -1.03*** | -1.02 | 1.00 |
| Task_157: Ximenic acid de novo synthesis | -1.04*** | -1.05*** | -1.04** | 1.00 |
| Task_158: Linolenate de novo synthesis | NA | NA | NA | NA |
| Task_159: Stearidonic acid de novo synthesis | -1.01 | -1.02** | 1.02 | 1.04*** |
| Task_160: omega-3-Arachidonic acid de novo synthesis | -1.01 | -1.02** | 1.00 | 1.02* |
| Task_161: EPA de novo synthesis | -1.01 | -1.03** | 1.02 | 1.04*** |
| Task_162: DPA de novo synthesis | -1.01 | -1.02** | 1.00 | 1.02* |
| Task_163: 9Z,12Z,15Z,18Z,21Z-TPA de novo synthesis | -1.01 | -1.02** | 1.00 | 1.02** |
| Task_164: 6Z,9Z,12Z,15Z,18Z,21Z-THA de novo synthesis | -1.01 | -1.02** | 1.00 | 1.02** |
| Task_165: DHA de novo synthesis | -1.01 | -1.02* | 1.01 | 1.03** |
| Task_166: 11Z,14Z,17Z-eicosatrienoic acid de novo synthesis | -1.02 | -1.03** | 1.00 | 1.02* |
| Task_167: 13,16,19-docosatrienoic acid de novo synthesis | -1.02 | -1.03** | 1.00 | 1.02* |
| Task_168: 10,13,16,19-docosatetraenoic acid de novo synthesis | -1.01 | -1.02** | 1.00 | 1.02* |
| Task_169: 12,15,18,21-tetracosatetraenoic acid de novo synthesis | -1.01 | -1.02** | 1.00 | 1.02* |
| Task_170: Linoleate de novo synthesis | NA | NA | NA | NA |
| Task_171: gamma-Linolenate de novo synthesis | -1.36*** | -1.44*** | -1.24** | 1.11 |
| Task_172: Dihomo-gamma-linolenate de novo synthesis | 1.01 | -1.02 | 1.05** | 1.07*** |
| Task_173: Arachidonate de novo synthesis | -1.03** | -1.04*** | -1.01 | 1.02** |
| Task_174: Adrenic acid de novo synthesis | -1.01 | -1.02** | 1.01 | 1.03** |
| Task_175: 9Z,12Z,15Z,18Z-TTA de novo synthesis | -1.01 | -1.02** | 1.01 | 1.03** |
| Task_176: 6Z,9Z,12Z,15Z,18Z-TPA de novo synthesis | -1.01 | -1.02** | 1.01 | 1.03** |
| Task_177: 4Z,7Z,10Z,13Z,16Z-DPA de novo synthesis | -1.01 | -1.02** | 1.00 | 1.02* |
| Task_178: 11Z,14Z-eicosadienoic acid de novo synthesis | -1.01 | -1.02** | 1.00 | 1.02* |
| Task_179: 13Z,16Z-docosadienoic acid de novo synthesis | -1.01 | -1.02** | 1.00 | 1.02* |
| Task_180: 10,13,16-docosatriynoic acid de novo synthesis | -1.01 | -1.02** | 1.01 | 1.02** |
| Task_181: Lauric acid complete oxidation | -1.05*** | -1.04*** | -1.05*** | -1.02 |
| Task_182: Tridecylic acid complete oxidation | -1.05*** | -1.04*** | -1.06*** | -1.03 |
| Task_183: Myristic acid complete oxidation | -1.05*** | -1.04*** | -1.05*** | -1.02 |
| Task_184: 9E-tetradecenoic acid complete oxidation | -1.05*** | -1.04*** | -1.06*** | -1.02 |
| Task_185: 7Z-tetradecenoic acid complete oxidation | -1.05*** | -1.04*** | -1.05*** | -1.02 |
| Task_186: Physeteric acid complete oxidation | -1.05*** | -1.04*** | -1.05*** | -1.02 |
| Task_187: Pentadecylic acid complete oxidation | -1.05*** | -1.04*** | -1.06*** | -1.02 |
| Task_188: Palmitate complete oxidation | -1.04*** | -1.04*** | -1.05*** | -1.02 |
| Task_189: Palmitolate complete oxidation | -1.04*** | -1.04*** | -1.05*** | -1.02 |
| Task_190: 7-palmitoleic acid complete oxidation | -1.04*** | -1.04*** | -1.05*** | -1.02 |
| Task_191: Margaric acid complete oxidation | -1.05*** | -1.04*** | -1.05*** | -1.02 |
| Task_192: 10Z-heptadecenoic acid complete oxidation | -1.05*** | -1.04*** | -1.05*** | -1.02 |
| Task_193: 9-heptadecylenic acid complete oxidation | -1.05*** | -1.04*** | -1.05*** | -1.02 |
| Task_194: Stearate complete oxidation | -1.04*** | -1.04*** | -1.05*** | -1.02 |
| Task_195: 13Z-octadecenoic acid complete oxidation | -1.04*** | -1.04*** | -1.05*** | -1.02 |
| Task_196: cis-vaccenic acid complete oxidation | -1.04*** | -1.04*** | -1.05*** | -1.02 |
| Task_197: Oleate complete oxidation | -1.04*** | -1.04*** | -1.05*** | -1.02 |
| Task_198: Elaidate complete oxidation | -1.04*** | -1.04*** | -1.05*** | -1.02 |
| Task_199: 7Z-octadecenoic acid complete oxidation | -1.04*** | -1.04*** | -1.05*** | -1.02 |
| Task_200: 6Z,9Z-octadecadienoic acid complete oxidation | -1.04*** | -1.04*** | -1.05*** | -1.02 |
| Task_201: Nonadecylic acid complete oxidation | -1.05*** | -1.04*** | -1.05*** | -1.02 |
| Task_202: Eicosanoate complete oxidation | -1.04*** | -1.04*** | -1.05*** | -1.02 |
| Task_203: 13Z-Eicosenoic acid complete oxidation | -1.04*** | -1.04*** | -1.05*** | -1.02 |
| Task_204: cis-gondoic acid complete oxidation | -1.04*** | -1.04*** | -1.05*** | -1.02 |
| Task_205: 9-Eicosenoic acid complete oxidation | -1.04*** | -1.04*** | -1.05*** | -1.02 |
| Task_206: 8,11-Eicosadienoic acid complete oxidation | -1.04*** | -1.04*** | -1.05*** | -1.02 |
| Task_207: Mead acid complete oxidation | -1.04*** | -1.04*** | -1.05*** | -1.02 |
| Task_208: Henicosanoic acid complete oxidation | -1.04*** | -1.04*** | -1.05*** | -1.01 |
| Task_209: Behenic acid complete oxidation | -1.04*** | -1.04*** | -1.05*** | -1.02 |
| Task_210: cis-erucic acid complete oxidation | -1.04*** | -1.04*** | -1.05*** | -1.02 |
| Task_211: cis-cetoleic acid complete oxidation | -1.04*** | -1.04*** | -1.05*** | -1.02 |
| Task_212: Tricosanoic acid complete oxidation | -1.05*** | -1.04*** | -1.06*** | -1.02 |
| Task_213: Lignocerate complete oxidation | -1.05*** | -1.04*** | -1.06*** | -1.02 |
| Task_214: Nervonic acid complete oxidation | -1.04*** | -1.04*** | -1.05*** | -1.02 |
| Task_215: Cerotic acid complete oxidation | -1.04*** | -1.04*** | -1.05*** | -1.02 |
| Task_216: Ximenic acid complete oxidation | -1.04*** | -1.04*** | -1.05*** | -1.02 |
| Task_217: Linolenate complete oxidation | -1.04*** | -1.04*** | -1.05*** | -1.02 |
| Task_218: Stearidonic acid complete oxidation | -1.04*** | -1.04*** | -1.05*** | -1.02 |
| Task_219: omega-3-Arachidonic acid complete oxidation | -1.04*** | -1.04*** | -1.04*** | -1.02 |
| Task_220: EPA complete oxidation | -1.04*** | -1.04*** | -1.05*** | -1.02 |
| Task_221: DPA complete oxidation | -1.04*** | -1.04*** | -1.05*** | -1.02 |
| Task_222: 9Z,12Z,15Z,18Z,21Z-TPA complete oxidation | -1.04*** | -1.04*** | -1.05*** | -1.02 |
| Task_223: 6Z,9Z,12Z,15Z,18Z,21Z-THA complete oxidation | -1.04*** | -1.04*** | -1.05*** | -1.02 |
| Task_224: DHA complete oxidation | -1.04*** | -1.04*** | -1.05*** | -1.02 |
| Task_225: 11Z,14Z,17Z-eicosatrienoic acid complete oxidation | -1.04*** | -1.04*** | -1.05*** | -1.02 |
| Task_226: 13,16,19-docosatrienoic acid complete oxidation | -1.04*** | -1.04*** | -1.05*** | -1.02 |
| Task_227: 10,13,16,19-docosatetraenoic acid complete oxidation | -1.04*** | -1.04*** | -1.05*** | -1.02 |
| Task_228: 12,15,18,21-tetracosatetraenoic acid complete oxidation | -1.05*** | -1.04*** | -1.06*** | -1.02 |
| Task_229: Linoleate complete oxidation | -1.04*** | -1.04*** | -1.05*** | -1.02 |
| Task_230: gamma-Linolenate complete oxidation | -1.04*** | -1.04*** | -1.05*** | -1.02 |
| Task_231: Dihomo-gamma-linolenate complete oxidation | -1.04*** | -1.04*** | -1.05*** | -1.02 |
| Task_232: Arachidonate complete oxidation | -1.04*** | -1.04*** | -1.05*** | -1.02 |
| Task_233: Adrenic acid complete oxidation | -1.04*** | -1.04*** | -1.04** | -1.01 |
| Task_234: 9Z,12Z,15Z,18Z-TTA complete oxidation | -1.04*** | -1.04*** | -1.04*** | -1.01 |
| Task_235: 6Z,9Z,12Z,15Z,18Z-TPA complete oxidation | -1.04*** | -1.04*** | -1.05*** | -1.02 |
| Task_236: 4Z,7Z,10Z,13Z,16Z-DPA complete oxidation | -1.04*** | -1.04*** | -1.05*** | -1.02 |
| Task_237: 11Z,14Z-eicosadienoic acid complete oxidation | -1.04*** | -1.04*** | -1.05*** | -1.02 |
| Task_238: 13Z,16Z-docosadienoic acid complete oxidation | -1.04*** | -1.04*** | -1.05*** | -1.03 |
| Task_239: 10,13,16-docosatrienoic acid complete oxidation | -1.04*** | -1.04*** | -1.05*** | -1.02 |
| Task_240: Triacylglycerol de novo synthesis | -1.04*** | -1.05*** | -1.02* | 1.03** |
| Task_241: Glycocholate de novo synthesis and excretion | -1.03* | -1.05*** | -1.00 | 1.03 |
| Task_242: Glycochenodeoxycholate de novo synthesis and excretion | -1.02* | -1.02* | -1.02 | -1.02 |
| Task_243: Taurocholate de novo synthesis and excretion | -1.03* | -1.05*** | -1.00 | 1.03 |
| Task_244: Taurochenodeoxycholate de novo synthesis and excretion | 1.03 | 1.00 | 1.07*** | 1.06*** |
| Task_245: PAPS de novo synthesis | -1.13*** | -1.11*** | -1.15*** | -1.06* |
| Task_246: PAPS degradation | -1.04*** | -1.05*** | -1.03** | 1.02 |
| Task_247: SAM de novo synthesis | -1.10*** | -1.05*** | -1.18*** | -1.12*** |
| Task_248: GSH de novo synthesis | -1.05*** | -1.04*** | -1.06*** | -1.04* |
| Task_249: Bilirubin conjugation | NA | NA | NA | NA |
| Task_250: NH3 import and degradation | -1.01 | -1.02** | 1.00 | 1.02* |
| Task_251: Ethanol import and degradation | -1.03** | -1.01 | -1.05** | -1.04** |
| Task_252: Oxidative phosphorylation | 1.07 | 1.02 | **1.16\*\*\*** | 1.07 |
| Task_253: Oxidative decarboxylation | **-1.47\*\*\*** | **-1.36\*\*\*** | **-1.68\*\*\*** | **-1.31\*\*\*** |
| Task_254: Krebs cycle NADH | 1.00 | 1.01 | -1.01 | -1.02 |
| Task_255: Ubiquinol-to-proton | 1.02 | 1.00 | 1.05* | 1.05** |
| Task_256: Ubiquinol-to-ATP | 1.01 | -1.00 | 1.03* | 1.04** |
| Task_257: Biosynthesis of Vitamin C | NA | NA | NA | NA |
| Task_259: Reactive oxygen species detoxification | -1.08** | -1.03 | **-1.18\*\*\*** | **-1.15\*\*\*** |
| Task_260: Presence of the thioredoxin system through the thioredoxin reductase activity | **-1.30\*\*\*** | **-1.35\*\*\*** | **-1.24\*\*\*** | 1.05 |
| Task_261: Inosine monophosphate synthesis | -1.10*** | -1.08*** | **-1.13\*\*\*** | -1.06*** |
| Task_262: GMP salvage from guanine | 1.03* | 1.04** | 1.01 | -1.03 |
| Task_263: ATP generation from ions | NA | NA | NA | NA |
| Task_264: Degradation of adenine to urate | **-1.11\*\*\*** | **-1.12\*\*\*** | **-1.11\*\*\*** | -1.01 |
| Task_265: Degradation of guanine to urate | -1.06*** | -1.06*** | -1.05* | 1.02 |
| Task_266: Degradation of cytosine | 1.01 | -1.01 | 1.06* | 1.06** |
| Task_267: Degradation of uracil | 1.01 | -1.01 | 1.06* | 1.06** |
| Task_268: Gluconeogenesis from pyruvate | -1.01 | -1.05*** | 1.05*** | 1.09*** |
| Task_269: Gluconeogenesis from Glutamine | -1.02 | 1.02 | -1.09*** | **-1.11\*\*\*** |
| Task_270: Glucose to lactate conversion | 1.00 | -1.02 | 1.04** | 1.06*** |
| Task_271: Malate to pyruvate conversion | **-1.11\*\*\*** | -1.03 | **-1.25\*\*\*** | **-1.23\*\*\*** |
| Task_272: Synthesis of erythrose-4-phosphate from fructose-6-phosphate | **-1.16\*\*\*** | **-1.21\*\*\*** | -1.10** | **1.12\*\*** |
| Task_273: Synthesis of ribose-5-phosphate | -1.10*** | -1.10*** | **-1.10\*\*\*** | -1.02 |
| Task_274: Synthesis of lactose | **-1.70\*\*\*** | **-1.48\*\*\*** | **-2.20\*\*\*** | **-1.93\*** |
| Task_275: Fructose to glucose conversion | **-1.10\*\*\*** | **-1.12\*\*\*** | -1.08** | 1.03 |
| Task_276: Mannose degradation | 1.06** | 1.03 | 1.10** | 1.06* |
| Task_278: Synthesis of acetone | -1.05*** | -1.06*** | -1.03* | 1.02 |
| Task_279: Synthesis of inositol | 1.02 | -1.04 | **1.11\*\*** | **1.13\*\*\*** |
| Task_280: Inositol as input for glucuronate-xylulose pathway | 1.04* | -1.01 | **1.12\*\*\*** | **1.13\*\*\*** |
| Task_281: Synthesis of phosphatidylinositol from inositol | **1.10\*\*\*** | 1.02 | **1.23\*\*\*** | **1.20\*\*\*** |
| Task_282: Conversion of 1-phosphatidyl-1D-myo-inositol 4,5-bisphosphate to 1D-myo-inositol 1,4,5-trisphosphate | **-1.23\*\*\*** | **-1.15\*\*\*** | **-1.37\*\*\*** | **-1.25\*\*\*** |
| Task_283: Starch degradation | **1.18\*\*** | **1.32\*\*\*** | -1.03 | **-1.28\*\*\*** |
| Task_284: Link between glyoxylate metabolism and pentose phosphate pathway | 1.06*** | 1.04** | 1.09*** | 1.04*** |
| Task_285: Synthesis of methylglyoxal | 1.02* | 1.02 | 1.03* | 1.01 |
| Task_286: Synthesis of alanine from glutamine | NA | NA | NA | NA |
| Task_287: Synthesis of arginine from glutamine | -1.03*** | -1.04*** | -1.03** | -1.01 |
| Task_288: Synthesis of nitric oxide from arginine | -1.08 | **-1.11\*** | -1.03 | 1.07 |
| Task_289: Synthesis of aspartate from glutamine | -1.00 | 1.00 | -1.01 | -1.01 |
| Task_290: Conversion of aspartate to arginine | **-1.12\*\*\*** | **-1.19\*\*\*** | -1.02 | **1.14\*\*\*** |
| Task_291: Conversion of aspartate to beta-alanine | 1.02*** | 1.02** | 1.03*** | -1.00 |
| Task_292: Conversion of asparatate to asparagine | -1.04 | **-1.11\*** | 1.06* | **1.16\*\*\*** |
| Task_293: Conversion of carnosine to beta-alanine | -1.03* | -1.07*** | 1.02 | 1.08*** |
| Task_294: Beta-alanine to 3-oxopropanoate | 1.04 | **1.14\*\*\*** | **-1.14\*** | **-1.32\*\*\*** |
| Task_295: Synthesis of cysteine from cystine | NA | NA | NA | NA |
| Task_296: Synthesis of taurine from cysteine | 1.08* | 1.03 | **1.16\*\*** | **1.11** |
| Task_297: Conversion of glutamate to glutamine | **-1.32\*\*\*** | **-1.30\*\*\*** | **-1.35\*\*\*** | -1.05 |
| Task_298: Conversion of glutamate to proline | **1.16\*\*\*** | **1.11\*\*\*** | **1.22\*\*\*** | 1.09*** |
| Task_299: Conversion of GABA into succinate | 1.09*** | **1.17\*\*\*** | -1.05 | **-1.25\*\*\*** |
| Task_300: Glutaminolysis | -1.02 | 1.02 | -1.08*** | **-1.10\*\*\*** |
| Task_301: Conversion of glycine to pyruvate | **-1.12\*\*\*** | **-1.18\*\*\*** | -1.04* | 1.09* |
| Task_302: Conversion of histidine to glutamate | -1.06*** | -1.06*** | -1.06*** | 1.00 |
| Task_303: Conversion of histidine to histamine | **1.50\*\*\*** | **1.44\*\*\*** | **1.58\*\*\*** | 1.10 |
| Task_304: Conversion of leucine to acetyl-coA | 1.01 | 1.05* | -1.05 | **-1.15\*\*\*** |
| Task_305: Conversion of lysine to L-Saccharopine | **-1.57\*\*\*** | **-1.32\*\*\*** | **-2.23\*\*\*** | **-1.60\*\*\*** |
| Task_306: Conversion of lysine to L-2-aminoadipate | **1.29\*\*\*** | **1.26\*\*\*** | **1.35\*\*\*** | 1.04 |
| Task_307: Synthesis of ornithine from glutamine | 1.04** | 1.08*** | -1.04 | **-1.13\*\*\*** |
| Task_308: Synthesis of spermidine from ornithine | **-1.10\*\*\*** | -1.08*** | **-1.14\*\*\*** | -1.06*** |
| Task_309: Phenylalanine to phenylacetaldehyde | -1.02* | 1.01 | -1.07*** | -1.06*** |
| Task_310: Phenylalanine to phenylacetyl-L-glutamate | -1.03** | -1.01* | -1.04** | -1.02* |
| Task_311: Phenylalanine to tyrosine | -1.00 | -1.00 | 1.00 | 1.00 |
| Task_312: Synthesis of proline from glutamine | 1.07*** | 1.10*** | 1.04 | -1.06** |
| Task_313: Synthesis of anthranilate from tryptophan | **1.19\*\*\*** | 1.04** | **1.41\*\*** | **1.40** |
| Task_314: Synthesis of kynate from tryptophan | 1.01 | -1.06 | **1.12** | **1.17\*** |
| Task_315: Synthesis of L-kynurenine from tryptophan | 1.09* | 1.02 | **1.21\*** | **1.19** |
| Task_316: Synthesis of N-formylanthranilate from tryptophan | -1.03* | **-1.14\*\*** | **1.13** | **1.30** |
| Task_317: Synthesis of quinolinate from tryptophan | **1.22\*\*\*** | **1.12\*\*\*** | **1.37\*\*\*** | **1.22\*\*\*** |
| Task_318: Synthesis of serotonin from tryptophan | -1.02 | 1.01 | -1.06** | -1.07** |
| Task_319: Tyrosine to adrenaline | **-1.14\*\*\*** | **-1.24\*\*\*** | -1.02 | **1.19\*\*\*** |
| Task_320: Tyrosine to dopamine | 1.00 | 1.00 | 1.01 | 1.00 |
| Task_321: Tyrosine to acetoacetate and fumarate | 1.03* | -1.03 | **1.12\*\*\*** | **1.14\*\*\*** |
| Task_322: Valine to succinyl-coA | -1.04* | 1.02 | **-1.15\*\*\*** | **-1.17\*\*\*** |
| Task_323: Hydroxymethylglutaryl-CoA synthesis | -1.05*** | -1.06*** | -1.03 | 1.03* |
| Task_324: Cholesterol synthesis | **-1.11\*\*\*** | **-1.12\*\*\*** | -1.09*** | 1.01 |
| Task_325: mevalonate synthesis | -1.08*** | -1.02 | **-1.21\*\*\*** | **-1.20\*\*\*** |
| Task_326: Glycerol-3-phosphate synthesis | **-1.22\*\*\*** | **-1.14\*\*\*** | **-1.35\*\*\*** | **-1.19\*\*\*** |
| Task_327: Synthesis of malonyl-coa | -1.05*** | -1.06*** | -1.04* | 1.02 |
| Task_328: Synthesis of palmitoyl-CoA | -1.05*** | -1.06*** | -1.04** | 1.02 |
| Task_329: Synthesis of thromboxane from arachidonate | **-1.18\*\*\*** | **-1.30\*\*\*** | -1.03 | **1.25\*\*\*** |
| Task_330: Synthesis of glucocerebroside | -1.02*** | -1.03*** | -1.01 | 1.01 |
| Task_331: Synthesis of globoside | -1.03*** | -1.04*** | -1.02* | 1.01 |
| Task_332: Synthesis of ubiquinone from tyrosine | -1.01 | 1.00 | -1.04 | -1.05 |
| Task_333: Synthesis of bilirubin | **-1.17\*\*\*** | **-1.25\*\*\*** | -1.08 | **1.17\*\*\*** |
| Task_334: Phosphatidyl-inositol to glucosaminyl-acylphosphatidylinositol | -1.05*** | -1.03** | -1.07*** | -1.04** |
| Task_335: Glucosaminyl-acylphosphatidylinositol to deacylated-glycophosphatidylinositol | -1.03* | -1.00 | -1.07*** | -1.09*** |
| Task_336: Degradation of n2m2nmasn | -1.01 | -1.07*** | 1.07* | **1.13\*\*\*** |
| Task_337: Biosynthesis of m4mpdol_U | -1.01 | -1.04*** | 1.04*** | 1.07*** |
| Task_338: Biosynthesis of g3m8masn | -1.03* | -1.03 | -1.04 | -1.02 |
| Task_339: Degradation of s2l2fn2m2masn | 1.03 | -1.05* | **1.14\*\*\*** | **1.20\*\*\*** |
| Task_340: Biosynthesis of core2 | -1.08*** | -1.09*** | -1.05* | 1.03 |
| Task_341: Biosynthesis of core4 | -1.04* | -1.08*** | 1.02 | **1.11\*\*** |
| Task_342: Biosynthesis of Tn_antigen | -1.07** | **-1.14\*\*\*** | 1.02 | **1.15** |
| Task_343: Keratan sulfate biosynthesis from O-glycan | -1.04* | -1.07*** | -1.01 | 1.07* |
| Task_344: Keratan sulfate biosynthesis from O-glycan | -1.04* | -1.07*** | -1.01 | 1.07* |
| Task_345: Keratan sulfate degradation | -1.00 | -1.02* | 1.02 | 1.04* |
| Task_346: Keratan sulfate biosynthesis from N-glycan | -1.02 | -1.06*** | 1.04* | 1.08*** |
| Task_347: AMP salvage from adenine | 1.00 | **-1.21\*\*\*** | **1.29\*\*\*** | **1.53\*\*\*** |
| Task_348: IMP salvage from hypoxanthine | 1.03* | 1.04** | 1.01 | -1.03 |
| Task_349: Carnosine synthesis from beta-alanine and histidine | **1.60\*\*\*** | **1.54\*\*\*** | **1.70\*\*\*** | **1.10** |
| Task_350: Calcium pumping to SR | **-1.23\*\*\*** | **-1.16\*\*\*** | **-1.37\*\*\*** | **-1.18\*\*\*** |
| Task_351: Calcium flux through RyR2 | -1.07** | 1.04 | **-1.29\*\*\*** | **-1.34\*\*\*** |
| Task_352: Contraction | **-1.12\*\*\*** | **-1.11\*\*** | **-1.14\*\*** | -1.05 |

**Supplementary figure 3:**
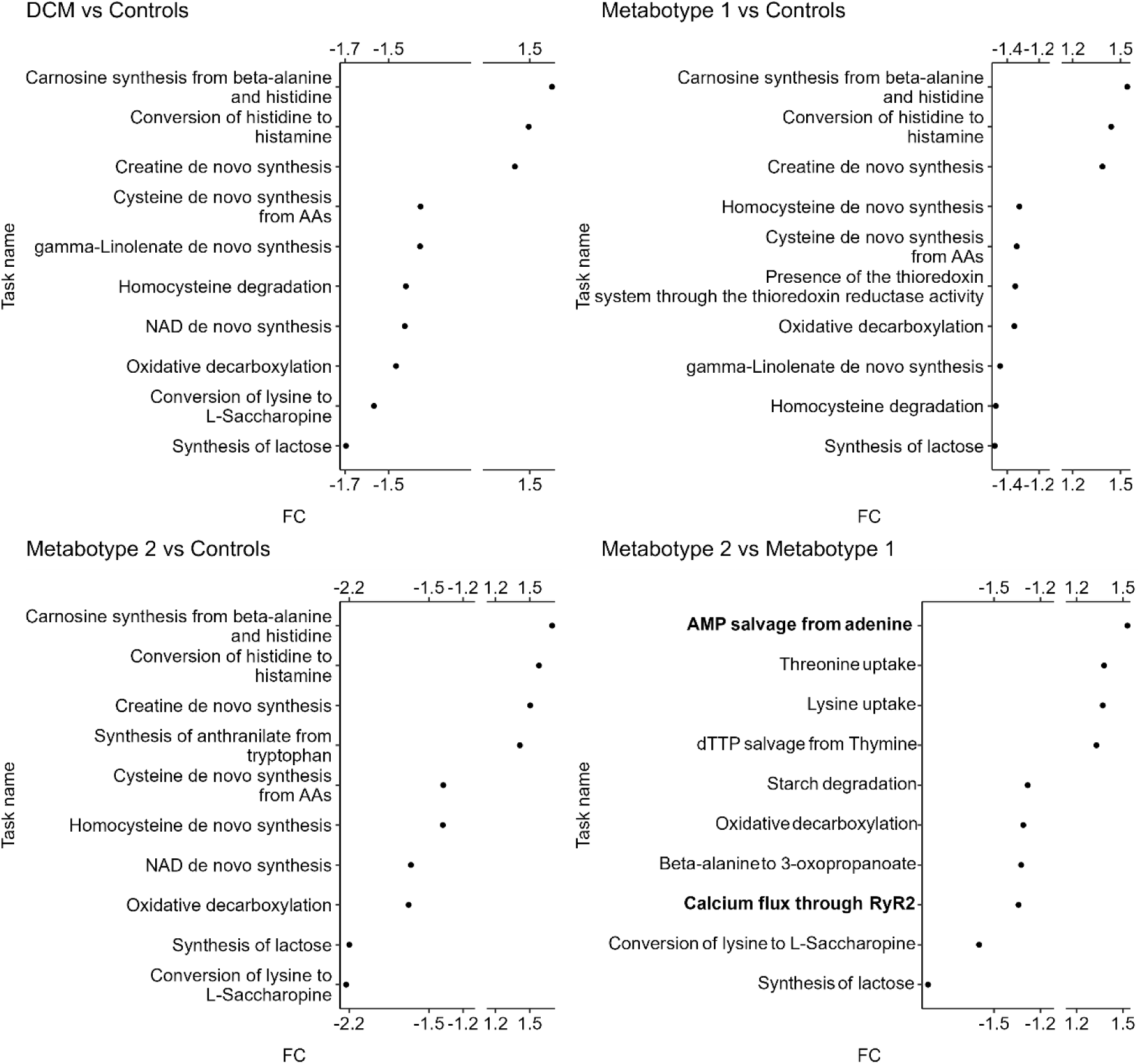
Fold-changes of the top 10 most different tasks (based on absolute fold-change) in each comparison made. FDR < 0.05 and absolute FC > 1.1 for all tasks presented. Task names in bold are part of the predefined tasks of interest. All of the tasks in the figure were verified to have sensible essential reactions, although task “Synthesis of anthranilate from tryptophan” only covers the conversion of tryptophan to L-formylkynurenine.

**Supplementary figure 4:**
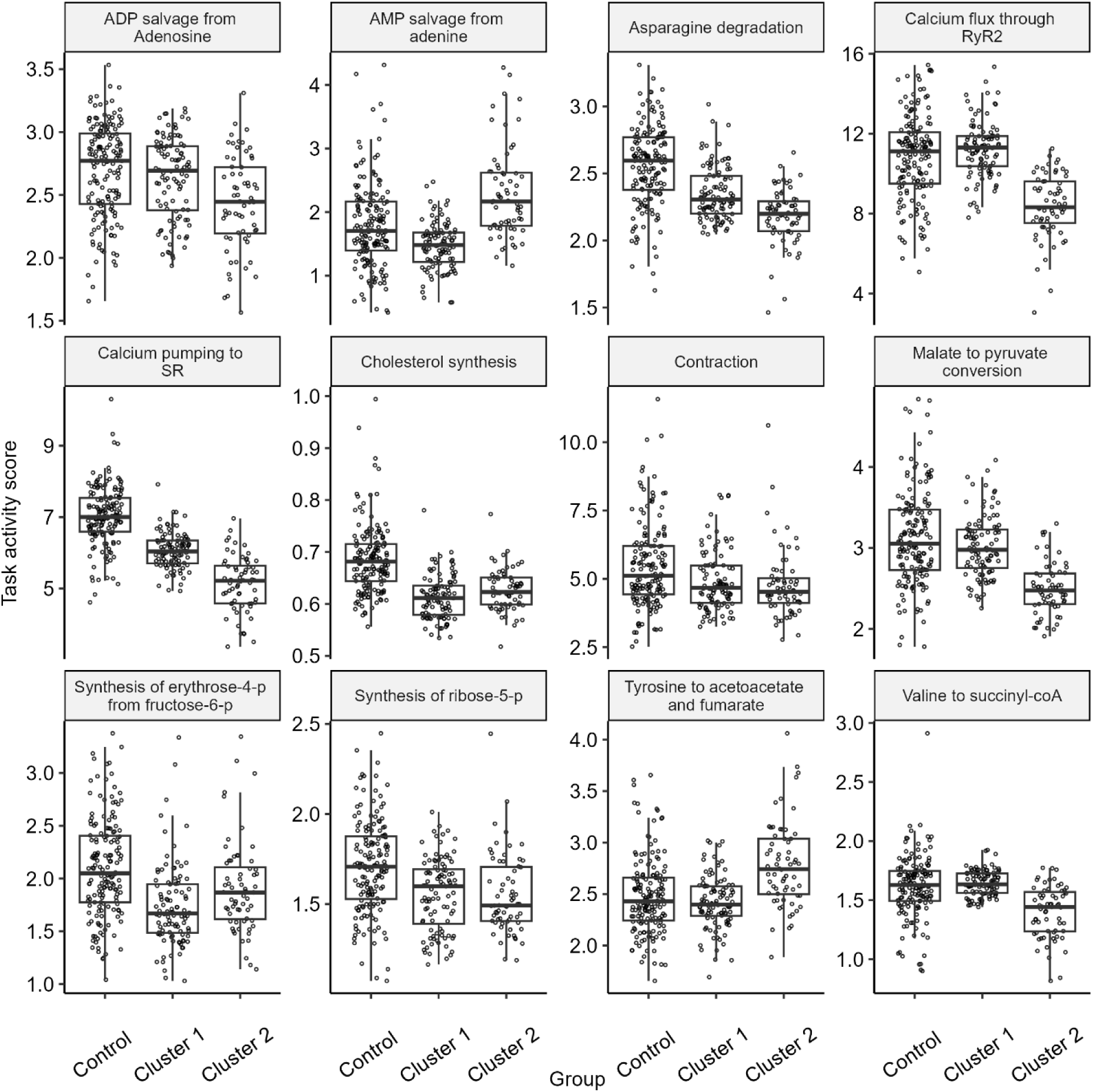
Boxplots showing task activity scores per group for each task of interest discussed in table 1.

**Supplementary figure 5:**
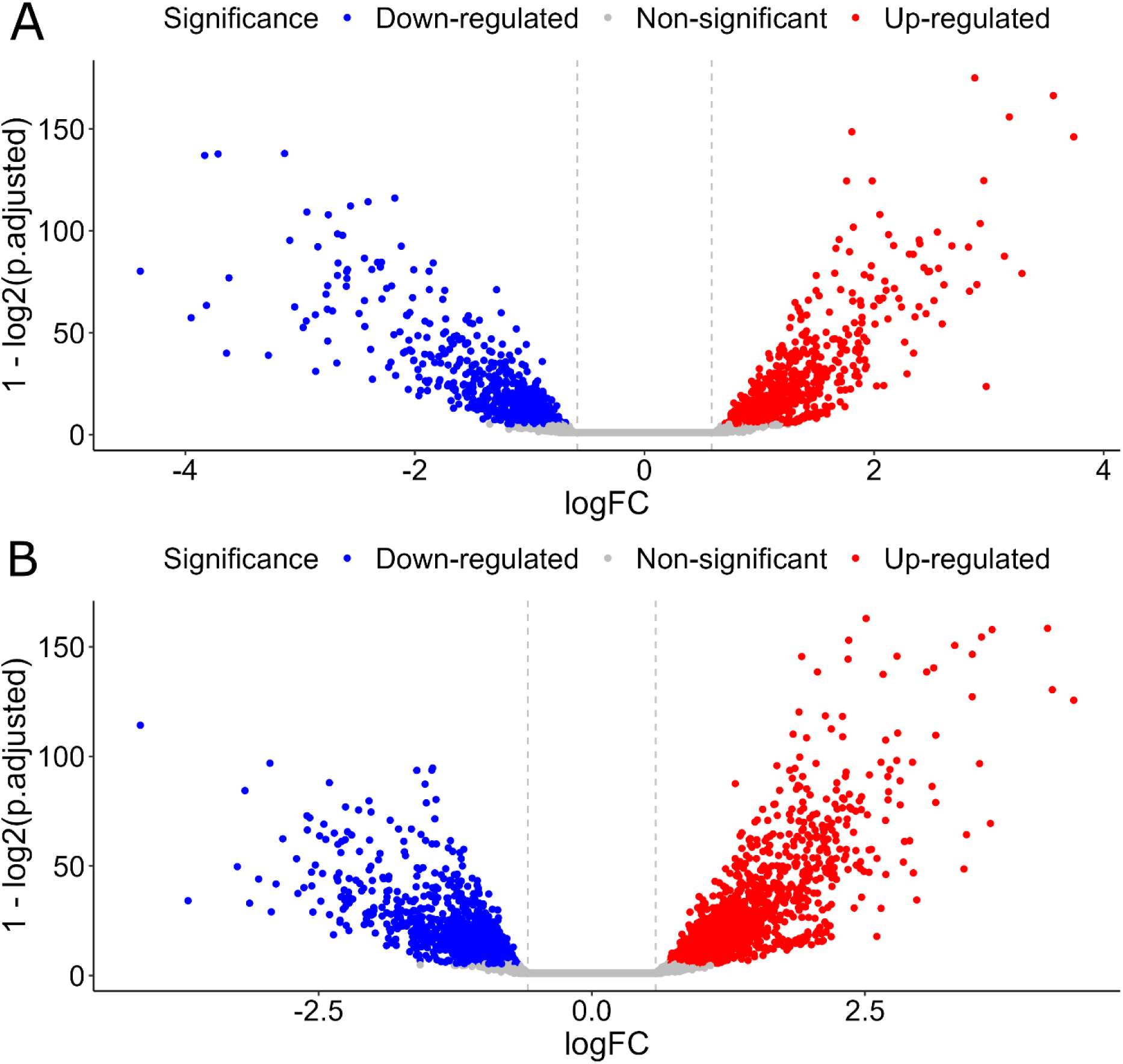
Volcano plot showing FDR-adjusted p-values in the differential expression analysis between controls and metabotypes 1 (A) and 2 (B).

**Supplementary figure 6:**
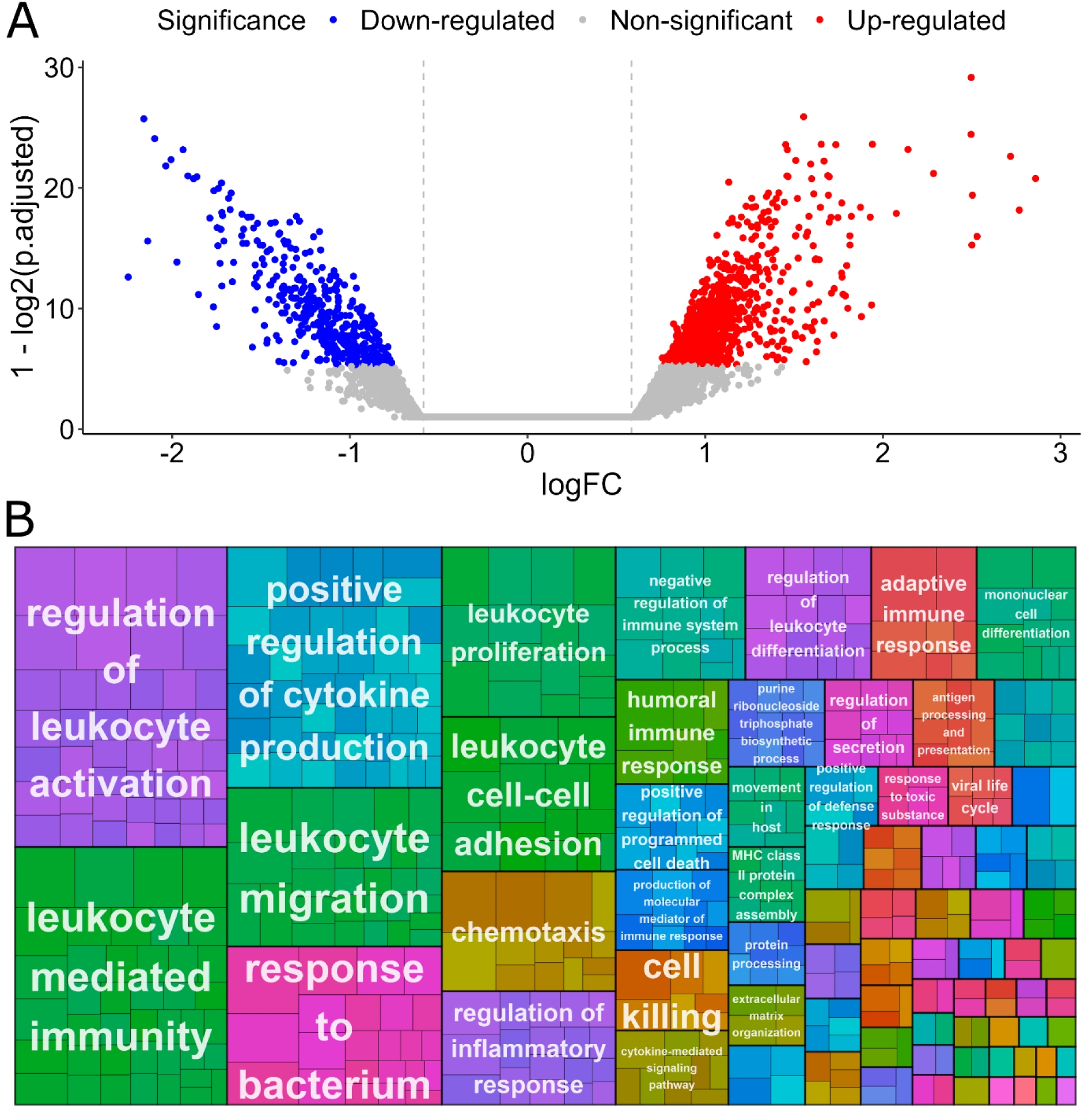
Volcano plot showing FDR-adjusted p-values in the differential expression analysis between metabotypes (A). Treemap plot of significantly enriched gene ontologies between metabotypes, grouped by similarity and represented by the most significant ontology in each group. Small rectangles represent individual ontologies with area relative to GSEA score. Colors assigned at random for visual contrast. (B).

**Supplementary figure 7:**
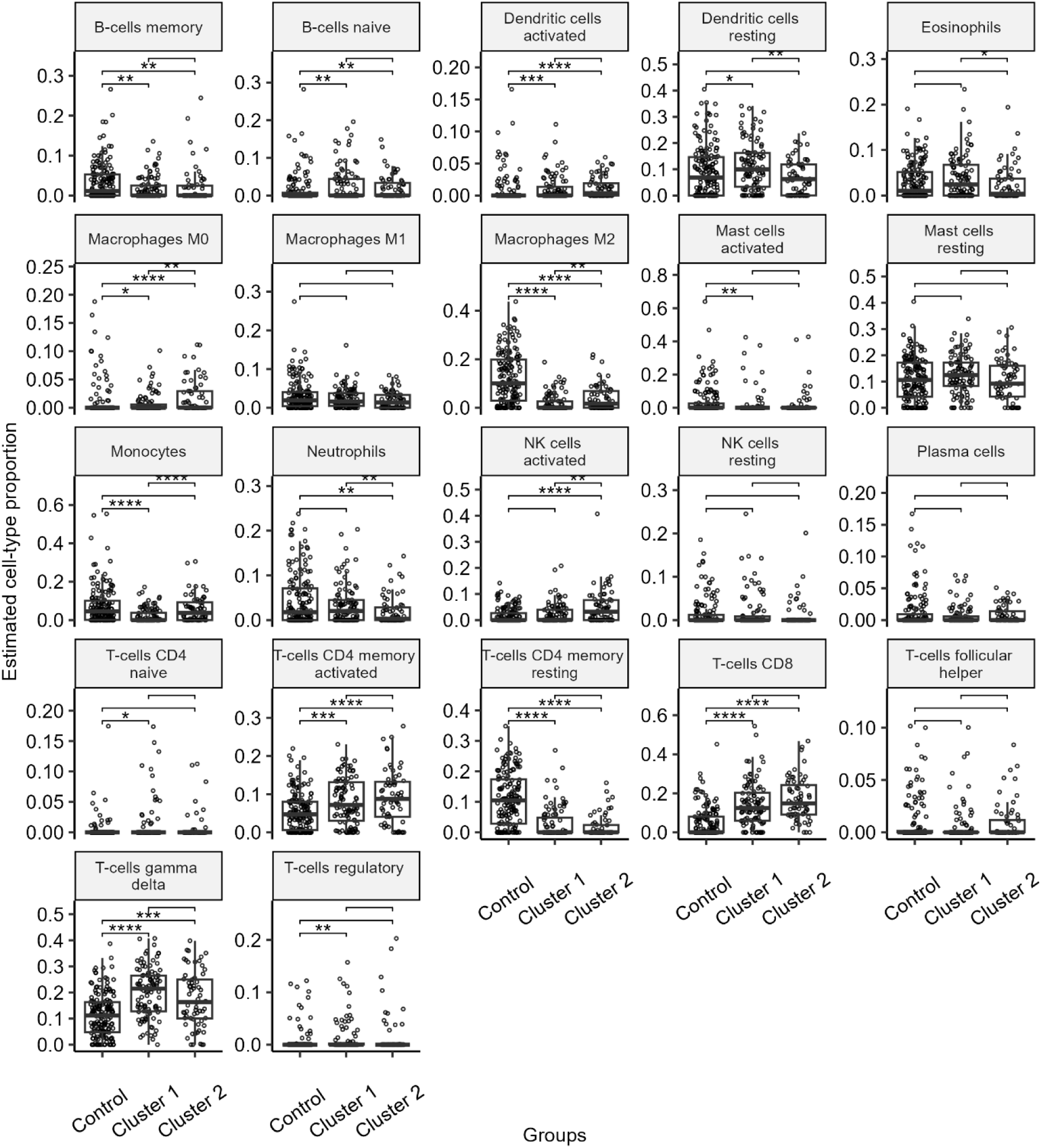
Boxplots showing estimated cell-type proportions imputed using CIBERSORT in the MaGNet dataset. * p-value < 0.05, ** p-value < 0.01, *** p-value < 0.001

